# Single-cell based integrative analysis of transcriptomics and genetics reveals robust associations and complexities for inflammatory diseases

**DOI:** 10.1101/2024.06.17.599349

**Authors:** Hope Townsend, Kaylee Rosenberger, Lauren Vanderlinden, Jun Inamo, Fan Zhang

## Abstract

**Background:** Understanding genetic underpinnings of immune-mediated inflammatory diseases is crucial to improve treatments. Single-cell RNA sequencing (scRNA-seq) identifies cell states expanded in disease, but often overlooks genetic causality due to cost and small genotyping cohorts. Conversely, large genome-wide association studies (GWAS) are commonly accessible.

**Methods:** We present a 3-step robust benchmarking analysis of integrating GWAS and scRNA-seq to identify genetically relevant cell states and genes in inflammatory diseases. <u>First</u>, we applied and compared the results of two recent algorithms, based on networks (scGWAS) or single-cell disease scores (scDRS), according to accuracy/sensitivity and interpretability (M. J. Zhang et al. 2022; Jia et al. 2022). While previous studies focused on coarse cell types, we used disease-specific, fine-grained single-cell atlases (183,742 and 228,211 cells) and GWAS data (Ns of 97,1,73 and 45,975) for rheumatoid arthritis (RA) and ulcerative colitis (UC) (F. Zhang et al. 2023; Ishigaki et al. 2022; Smillie et al. 2019; de Lange et al. 2017). <u>Second</u>, given the lack of scRNA-seq for many diseases with GWAS, we further tested the tools’ resolution limits by differentiating between similar diseases with only one fine-grained scRNA-seq atlas. <u>Lastly</u>, we provide a novel evaluation of noncoding SNP incorporation methods by testing which enabled the highest sensitivity/accuracy of known cell-state calls.

**Results:** We first found that single-cell based tool scDRS called superior numbers of supported cell states, like MERTK+ myeloid cells in RA, which were overlooked by network-based scGWAS. While scGWAS was advantageous for gene exploration, scDRS captured cellular heterogeneity of disease-relevance without single-cell genotyping. For noncoding SNP integration, we found a key trade-off between statistical power and confidence with positional (e.g. MAGMA) and non-positional approaches (e.g. chromatin-interaction, eQTL). Even when directly incorporating noncoding SNPs through 5’ scRNA-seq measures of regulatory elements, non disease-specific atlases gave misleading results by not containing disease-tissue specific transcriptomic patterns. Despite this criticality of tissue-specific scRNA-seq, we showed that scDRS enabled deconvolution of two similar diseases with a single fine-grained scRNA-seq atlas and separate GWAS. Indeed, we identified supported and novel genetic-phenotype linkages separating RA and ankylosing spondylitis, and UC and crohn’s disease. Overall, while noting evolving single-cell technologies, our study provides key findings for integrating expanding fine-grained scRNA-seq, GWAS, and noncoding SNP resources to unravel the complexities of inflammatory diseases.

## 1 Introduction

The efficacy of treatments for immune-mediated inflammatory diseases, such as rheumatoid arthritis (RA) and ulcerative colitis (UC), varies across patients(Morgan et al. 2003). Single-cell RNA sequencing (scRNA-seq) technology enables the development of effective treatments for patients with with immune-mediated inflammatory diseases by allowing the identification of specific cell states expanded in diseased tissue or blood(“Method of the Year 2019: Single-Cell Multimodal Omics” 2020). However, most scRNA-seq analyses do not consider genetic causality, and due to its high expense, available single cell datasets are often confined to small patient cohorts. Understanding the genetic underpinnings of diseases is key for preventative care, unraveling physiological and environmental contributions to pathology, and allowing personalized treatments. Genome wide association studies (GWAS) have been the gold standard to identify disease-associated genetic *loci* and summary statistics for large cohorts are often publicly accessible (Schaid, Chen, and Larson 2018). Therefore, recent work has gone into combining the physiological insights from scRNA-seq with genetic associations from GWAS for unraveling disease causality. Indeed attempts to integrate bulk RNA-seq studies with GWAS have been implemented, yet still only explain about 30% of the heritability by gene expression for complex traits(Yao et al. 2020). This pitfall is likely explained by the less fine-scale cell states available with bulk RNA-seq compared to scRNA-seq, where immune cells exhibit divergent expression profiles at nuanced cell states, with different cell phenotypes being uniquely associated with disease(Jew et al. 2020; Rao et al. 2017; F. Zhang et al. 2023).

Recently, several computational tools have been developed to link disease relevant *loci* from GWAS to nuanced cell states revealed by scRNA-seq to identify disease-associated cell states and the relevant genetics active in them(Timshel, Thompson, and Pers 2020; R. Wang, Lin, and Jiang 2022; Rouhana et al. 2021; M. J. Zhang et al. 2022; Jia et al. 2022). Major steps include first summarizing highly expressed genes from single cell expression data, linking GWAS based single nucleotide polymorphisms (SNPs) to genes in the scRNA-seq data, and then using statistical tests to identify significant associations. However, a thorough comparison and assessment of these tools is lacking. Notably, linking SNPs from GWAS to the genes they potentially impact has been notoriously difficult with no clear solution (Leeuw et al. 2015; Watanabe et al. 2017; Gazal et al. 2022; Weeks et al. 2023; Yang, Chen, and Zhao 2021). With more than 90% of immune-disease associated SNPs falling into noncoding regions, most of which are in cis-regulatory regions, the need to link these SNPs to genes cannot be overstated (Chen et al. 2020). For example, MAGMA links SNPs to genes according to a user-selected window size outside the gene (Leeuw et al. 2015). MAGMA can take both genotype data and summary statistics as input while accounting for Linkage Disequilibrium. It outputs a list of thousands of genes with the corresponding GWAS statistics reestablished at the gene level. However, many target genes of cis-regulatory regions are not the closest gene and can even be farther than 1 Mb away, contradicting the assumptions of tools like MAGMA. Therefore, alternative methods focusing on eQTL, chromatin contact (e.g. Hi-C), and similarly relevant enhancer-gene linking data have been introduced(Watanabe et al. 2017; Gazal et al. 2022). Indeed, newer studies have begun introducing single-cell transcriptomics methods that measure cis-regulatory elements to directly consider noncoding SNPs(Moody et al. 2023). The influence of incorporating noncoding SNPs using non-positional compared with positional methods, specifically within the context of these algorithms, has not been formally evaluated.

These algorithms have been previously applied to scRNA-seq atlases not specific to diseases, limiting results to capture only general associations (e.g. metabolic cells for metabolic diseases)(M. J. Zhang et al. 2022; Jia et al. 2022). Within single-cell technologies having revolutionized exploration of diseased tissue heterogeneity, many disease-specific atlases are still being generated. Thus, integrating genetics specific to under-studied diseases with in-depth scRNA-seq cell annotations for the same or a clinically similar disease might enable the identification of significant and novel cell states.

Overall, despite the recent influx of tools integrating genetics and single cell transcriptomics, a thorough comparison and assessment of different types of recent algorithms and major challenges of the domain is lacking. To address this, we benchmarked representative algorithms, scGWAS and scDRS, by unbiasedly linking GWAS with single-cell phenotypes targeted on four immune-mediated disease datasets (F. Zhang et al. 2023; M. J. Zhang et al. 2022; Jia et al. 2022; Smillie et al. 2019). We further annotated our results based on literature support of calls, and evaluated the computational efficiency and result interpretability. Given most immune relevant SNPs are noncoding, we then evaluated the influence of different methods incorporating these SNPs for use in the algorithms (Leeuw et al. 2015; Watanabe et al. 2017). As a result, we first showed that both scGWAS and scDRS successfully identified expected significant cell types for tested diseases when using fine-grained scRNA-seq atlases, although scDRS identified more significant results with literary support. We also found that scDRS can be used to distinguish cell phenotypes for different diseases while using the same fine-grained scRNA-seq atlas. Finally, we provided evidence supporting the usage of positional based methods to incorporate noncoding SNPs until other methods can increase in statistical power and use more relevant atlases. Overall, our in-depth benchmarking and application on disease tissue data demonstrated that current tools could identify associations between cell phenotypes and disease with high resolution and specificity, and also pointed out the lack of in-depth atlases requiring future improvement.

## 2 Methods & Materials

**Figure 1.**
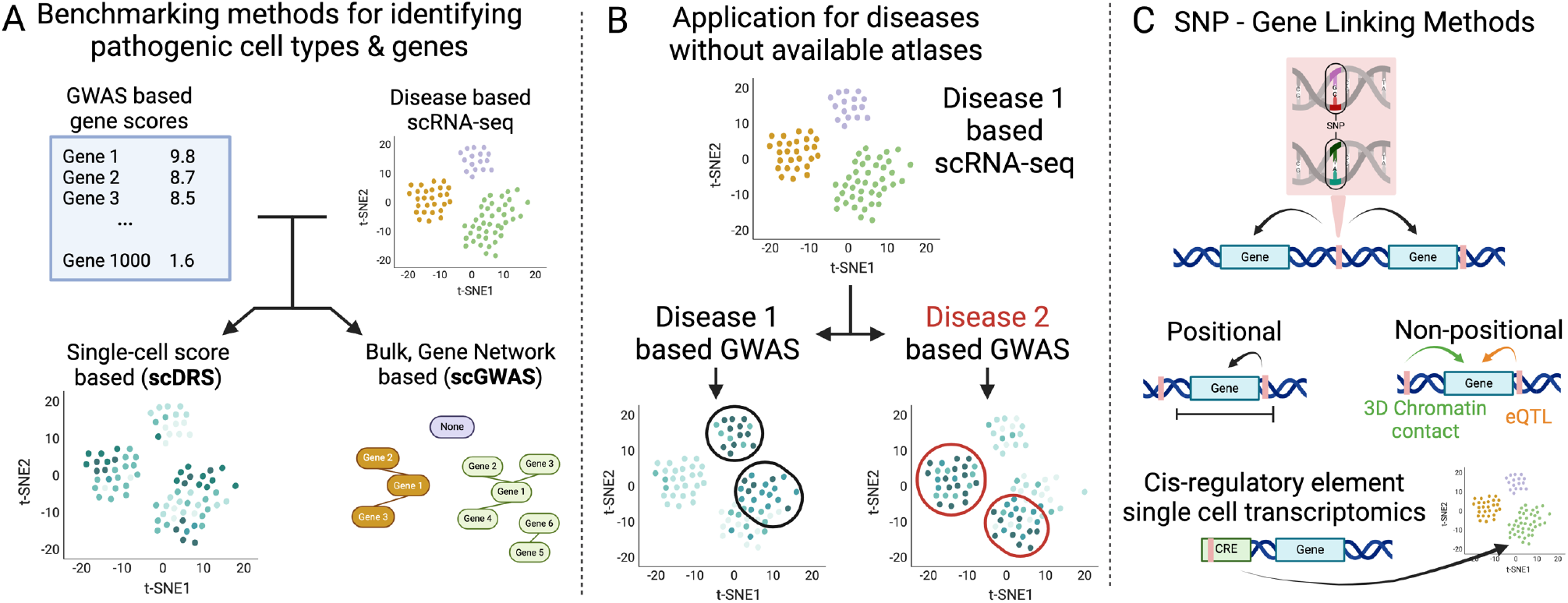
Overview of study design. We first benchmarked the two most recent and representative algorithms in the field according to the number of literature supported clusters called significant, computational efficiency, and result interpretability (Figure 1A). Expected results were based on a literature search for each individual cell phenotype for expansion in a disease and/or genetic connections, the results of which can be found in Supplemental Tables 1 and 2. If a cell type with multiple cell states was significant, the cell states were marked as having “general” literature support while if a specific cell state was supported, it had “specific” literature support. Due to the robustness of the available atlases and studies, we used scRNA-seq data generated from inflamed RA synovial and UC colon to determine disease-associated cell states (F. Zhang et al. 2023; Smillie et al. 2019). Next, we assessed the feasibility of using identical scRNA-seq atlases to distinguish between two clinically similar diseases, using RA inflamed synovial tissue for RA and ankylosing spondylitis (AS), and UC colon for UC and crohn’s disease (CD) (Figure 1B). Finally, we evaluated the incorporation of noncoding SNPs when using positional (MAGMA) vs non-positional based SNP-gene linking methods or cis-regulatory element focused single-cell omics like ATAC-seq or 5’-scRNA-seq (Figure 1C). We deploy all the code and analytical pipelines at our Github repository for reproducible research at https://github.com/fanzhanglab/SCRNA-GWAS-Benchmarking.

### 2.1 Selection of tools

We summarized the attributes of five currently-usable packages that integrate scRNA-seq data and GWAS summary statistics to identify significant cell types and/or the GWAS-linked genes that best explain these cell types (Table 1). Other methods are available such as RolyPoly, CocoNet, and sc-linker, but appear to be no longer supported or do not provide all necessary code as the methods were not designed as user-friendly packages(Calderon et al. 2017; Jagadeesh et al. 2022; Shang, Smith, and Zhou 2020). Single-cell disease score generation (scDRS) and gene network comparison (scGWAS) methods were the most recent tools and provide unique results as either gene networks and single-cell based results. Other methods differ most by their incorporation of noncoding SNPs which is addressed separately in this work.

**Table 1.** Summary table of the currently available and operable tools for identifying significant cell types and/or genes based on the integration of GWAS and single-cell RNA-seq data. Non-operable methods (RolyPoly) and methods that do not include all necessary code to run the entire pipeline (CocoNet and SC-Linker) are not included in the table.

| Package (Year) Interface | Inputs | Outputs | SNP-Gene Linking | Summary | Highlights |
| --- | --- | --- | --- | --- | --- |
| **scGWAS** (2022) CL JAR, pre/post processing in R | 1.Boxcox transformed gene p-values<br>2. Pseudobulk<br>3. Gene-gene network file | 1. Significant gene modules in each cell type | Window-based: MAGMA | Network-based approach to identify cell types overexpressed with disease-significant genes | Pathway based for more meaningful output |
| **scDRS** (2022) CLI or API | 1.Anndata single cell expression data<br>2.Gene p-values or z-scores | 1.Cell score file for a given trait<br>2. Opt: Cell group score and heterogeneity<br>3.Opt: Cell variable (e.g. gene) correlation to disease scores | Window-based: MAGMA | Monte Carlo simulation method that scores individual cells for disease association based on increased expression of sets of putative disease genes | Single-cell level allows unique post analyses |
| **EPIC**<br>(2022)<br>R package | 1.Pseudobulk gene expression<br>2.GWAS summary stats | 1.Enrichment score of trait for each cell type<br>2.Relevant genes from DFBETAS | Sliding-window based LDSC | Gene-level chi-square association testing, then gene-level regression-association testing for each cell type | Adapted for rare and common variants |
| **ECLIPSER**<br>(2021)<br>Scripts on Github | 1.GWAS summary stats<br>2.Gene differential expression table | 1.Prioritized cell types<br>2.Leading edge causal genes and eQTL impact | eQTL and other functional evidence | Cell-type specificity score for each GWAS locus, cell-type specific genes (from differential expression analysis mapped to locus) | Provides putative regulatory impact of genes |
| **CELLECT**<br>(2020)<br>CLI | 1.Specificity input from CELLEX<br>2.GWAS summary stats | 1.Prioritized cell types<br>2.Opt: Gene heritability | LDSC or MAGMA | Heritability regression based method with CELLEX gene specificity scores | Allows easy usage of LDSC or MAGMA |

### 2.2 Data availability

The GWAS data used in this work can be found in Supplemental Table 3. Due to the most robust LD score data belonging to those with European descent, and the larger sample size of this group in both GWAS and scRNA-seq data, we focused on this subpopulation for the purpose of this benchmarking analysis. The major histocompatibility complex region was not included due to its complex genetic architecture. For GWAS summary statistics without rsids for RA, SNPs were assigned to rsids using BEDOPs duplicate/synonymous rsids with the lowest p-values were kept. The code for these steps can be found on our github under SCRNA-GWAS-Benchmarking/src/00B_Preprocess_GWAS.

For RA and AS, we analyzed a scRNA-seq data set developed by Zhang et al., 2023. To stay consistent with GWAS data, we only included cells from individuals of European descent with RA, leaving 183,742 cells. We used their most updated cell-state and cell-type annotations determined by their analysis of 314,011 cells with transcriptomic, CITE-seq, experimental evidence and batch control to ensure the best validation. All expression was normalized with log(1 + UMIs for gene/totl UMIs in cell *10,000), and cells expressing fewer than 500 genes or that contained more than 20% if their total UMIs mapping to mitochondrial genes were removed. Further QC analysis is described in their paper(F. Zhang et al. 2023). For UC and CD, we analyzed the scRNA-seq dataset from Smillie et al., 2019 which contained 228,211 cells passing quality control. The code for these steps can be found on our github under SCRNA-GWAS-Benchmarking/src/00A_Preprocess_scRNA.

### 2.3 SNP-Gene Linking

MAGMA-based SNP-gene linking was done using version v1.10 with NCBI37.3.gene.loc and NCBI38.gene.loc downloaded from the MAGMA website as the gene locations files, and European UK Biobank Phase 3 LD scores. The window size of 50-35kb was chosen for the final comparison of significant cell clusters as it was the window size used in the original scGWAS paper and scDRS showed the most consistency with window size. When assessing the impact of this window size parameter on scDRS, sizes 0kb, 5kb, 10kb, and 100kb were also chosen based on the window sizes used across the literature (Supplemental Table 4). For this parameter stability assessment, only the top 1000 ranking genes according to MAGMA that were also found in the scRNA-seq expression data were used, so that a final total of 1000 genes were used. Synonyms according to genecards.org and humanproteinatlas.com and were also considered to verify proper comparison of genes between MAGMA and scRNA-seq. These databases were chosen over available NCBI tables as some needed synonyms were unique to these databases. If genes from the scRNA-seq dataset were still not found in the MAGMA file, they were added to allow their inclusion in the analysis. The genes identified by MAGMA but not found in scRNA-seq data are discussed further in the Supplemental Material, with numbers dictated in Supplemental Table 5.

The code for these steps can be found on our github under SCRNA-GWAS-Benchmarking/src/01_MAGMA_Gene_Alias.

FUMA is a web-based tool that determines statistically significant disease associated genes using positional, eQTL, and 3D chromatin based mapping, but does not calculate a summary p-value like MAGMA(Watanabe et al. 2017). Therefore, to explore the implications of including these forms of mapping, we used the minimum GWAS SNP P-value (minGwasP in genes.txt output file) for each gene as a proxy for a disease-association p-value to allow input for scDRS and scGWAS. GWAS input for FUMA was the unedited version of GWAS summary statistics as FUMA identifies lead SNPs, maps to rsIDs, addresses duplicate and synonymous rsIDs, and filters out the MHC region in its analysis. Default parameters were used including a MAGMA window of 10kb, with MAGMA expression data being based on GETx v8. We also used eQTL and Chromatin Interaction Mapping, both including the options of available blood cell eQTL data. Versions include FUMA v1.5.3, MAGMA v1.08, GWAScatalog e0_r2022-11-29, and ANNOVAR 2017-07-17.

### 2.4 scGWAS & scDRS

scGWAS uses a network-based approach to uncover cell types that significantly express disease-associated genes and identify gene modules representing disease-specific processes(Jia et al. 2022). Unlike other methods where cell types are assigned a disease-significance score, scGWAS assigns significance scores to gene modules with strong representation in both scRNA-seq cell type expression and GWAS based on a proportional test. scGWAS is implemented in Java via a JAR package (ver. scGWAS_r1.jar) on the author’s GitHub repository and can be run through the command line. To run the package, a configure file is required, which provides all input data and parameters for running scGWAS. Based on author recommendations on their GitHub repository, parameters were kept at default values. Further, we used the same PathwayCommons input network file as Jia and Zhao (2022), with gene-gene relationship information used for constructing the background network. We followed the analysis pipeline described on the authors’ GitHub repository for the following steps. For the screen expression input file, we processed the scRNA-seq dataset using their R-script to calculate the average log-transformed gene-based CPM per defined cell type. We processed the MAGMA output using the box-cox transformation script as the GWAS node input file. We ran scGWAS on the same scRNA-seq dataset first with general cell types and then on fine-scale defined clusters. The code for these steps can be found on our github under SCRNA-GWAS-Benchmarking/src/03_scGWAS.

scDRS is the only tool we identified that assesses disease-association at the individual cell level which also allows it to consider cell level covariates(M. J. Zhang et al. 2022). It then presents downstream analyses that use unified Monte Carlo tests to identify significant pre-annotated cell states according to a group Z score and genes whose expressions correlate with disease scores. The CLI version of scDRS was used according to their GitHub repository (https://github.com/martinjzhang/scDRS). Version v102 v1.0.2 was used according to their installation instructions. All default parameter values were used, and P-value files output from MAGMA served as input to scdrs munge-gs. The covariates files used in computing scDRS scores included nUMI, number of genes, and sex for both RA & UC, and age and duration for RA, and sample location, percent of mitochondrial reads, and smoking status for UC (found in our github at SCRNA-GWAS-Benchmarking/data/SC_data). We ran downstream analyses to identify on the same scRNA-seq dataset using annotations of general cell types and then with fine-scale defined clusters. The code for these steps can be found on our github under SCRNA-GWAS-Benchmarking/src/02_scDRS.

### 2.5 Benchmarking methods

Both packages provide results indicating which cell clusters are significant for the disease, but the exact format and calculation of these results differs. scGWAS provides significance in the form of gene modules within clusters that have disease-relevance, whereas scDRS provides single cell disease scores as well as Z-scores for annotated clusters and measurements regarding the heterogeneity of these disease scores within each cluster. To compare results across the two packages, we defined significant cell clusters in scGWAS as clusters with at least one disease-significant gene module. We then assessed whether the packages identified significant cell types similarly across a given disease.

For scDRS gene outputs, https://www.gsea-msigdb.org/gsea/msigdb/human/compute_overlap was used to establish what processes the top most positively correlated genes most overlapped with(Mootha et al. 2003; Subramanian et al. 2005). The following gene set collections were included: Hallmark, Curated (C2), Regulatory (C3), Biological Process (GOBP), and IMMUNESIGDB (C7-IMMUNE). The Cell type (C8) collection was not included to allow a more basic physiological understanding of results. For scGWAS, we completed a similar gene set overlap analysis for each significant gene module separately. Here, we included the cell type signature gene sets (C8) to ensure logical results given the smaller gene numbers.

## 3 Results

### 3.1 Benchmarking network-based and single-cell based methods reveals consistent associations of nuanced cell phenotypes with inflammatory diseases

We built our initial benchmarking pipeline on evaluating both cell types and finer grained cell states as well as gene modules on RA and UC datasets. Despite previous applications being largely confined to large scale cell types, both scDRS and scGWAS did not call all the cell types with significant fine-grained cell states significant (Supplemental Figure 1). scGWAS and scDRS mostly showed agreement in significantly called fine-grained cell states, although scDRS called far more significant clusters with literature support for connection with the disease (**Figure 2**). For example, scDRS uniquely identified two cell groups heavily implicated in RA: MERTK+ myeloid cells and a rare B-cell cluster called autoimmune-associated B-cells (ABCs). Ours and others work recently found that these two cell phenotypes are expanded in inflamed synovial tissues for RA, but the genetic association was not clear(F. Zhang et al. 2023; Kuo et al. 2019; Y. Wang et al. 2019). While the number of significant NK cell clusters in RA were comparable between scGWAS and scDRS (8 and 5), only 3 of the significantly called clusters were shared. Indeed the top scored clusters were opposite ranked between the two methods according to scDRS group Z-score for and the number of significant scGWAS clusters. Still, both called collections of CD56bright and CD56dim NK cells significant. CD56dim cells have decreased cytotoxicity in RA while CD56bright cells are expanded in inflamed joints, increasing the production of pro-inflammatory cytokines(Chalan et al. 2016; Dalbeth and Callan 2002; Yamin et al. 2019). Similarly, both methods called the IL7R+CD151+ ILC (NK-13) cells significant but not IL7R+ ILC (NK-12) cells, possibly clarifying the role of IL7R+ in NK cells that has been previously supported for RA(Meyer, Parmar, and Shahrara 2022; Churchman and Ponchel 2008). The lack of conservation for NK cell calls might be explained by the lack of study of NK cells in RA making relevant cluster separation more difficult. Importantly, neither algorithm identified significant fibroblast cell types despite NOTCH3+ and CD34+ sublining fibroblasts being expanded in RA, possibly indicating that these phenotypes are due to upstream genetic linkages as previously predicted(Phillips 2021). The small number of significant calls for UC is consistent with previous studies with scDRS. Yet, both scDRS and scGWAS identified a recently discovered cell group with the highest expression of putative IBD risk genes in inflamed vs healthy tissue corroborated by two separate cohorts (M cells)(Smillie et al. 2019; Serigado et al. 2022). Interestingly, neither algorithm called CD8+ IL17+ T cells or Tuft cells despite these groups having significantly different portions in UC(Smillie et al. 2019; Kjærgaard et al. 2021). The calling of T-regulatory cells and epithelial cells maintaining epithelial barrier integrity might instead point to the genetic casualty of these differing proportions, as transcriptional differences in both groups have been pointed to proportional shifts of cell states relevant to UC(Chelakkot, Ghim, and Ryu 2018; Rath and Haller 2022; Yamada et al. 2016). Finally, the median of individual scDRS disease Z-scores in cell clusters does not always correspond with the scDRS group Z-score of the clusters; this is most obvious in the NK cells from RA and Epithelial cells of UC (Figure 2).

**Figure 2.**
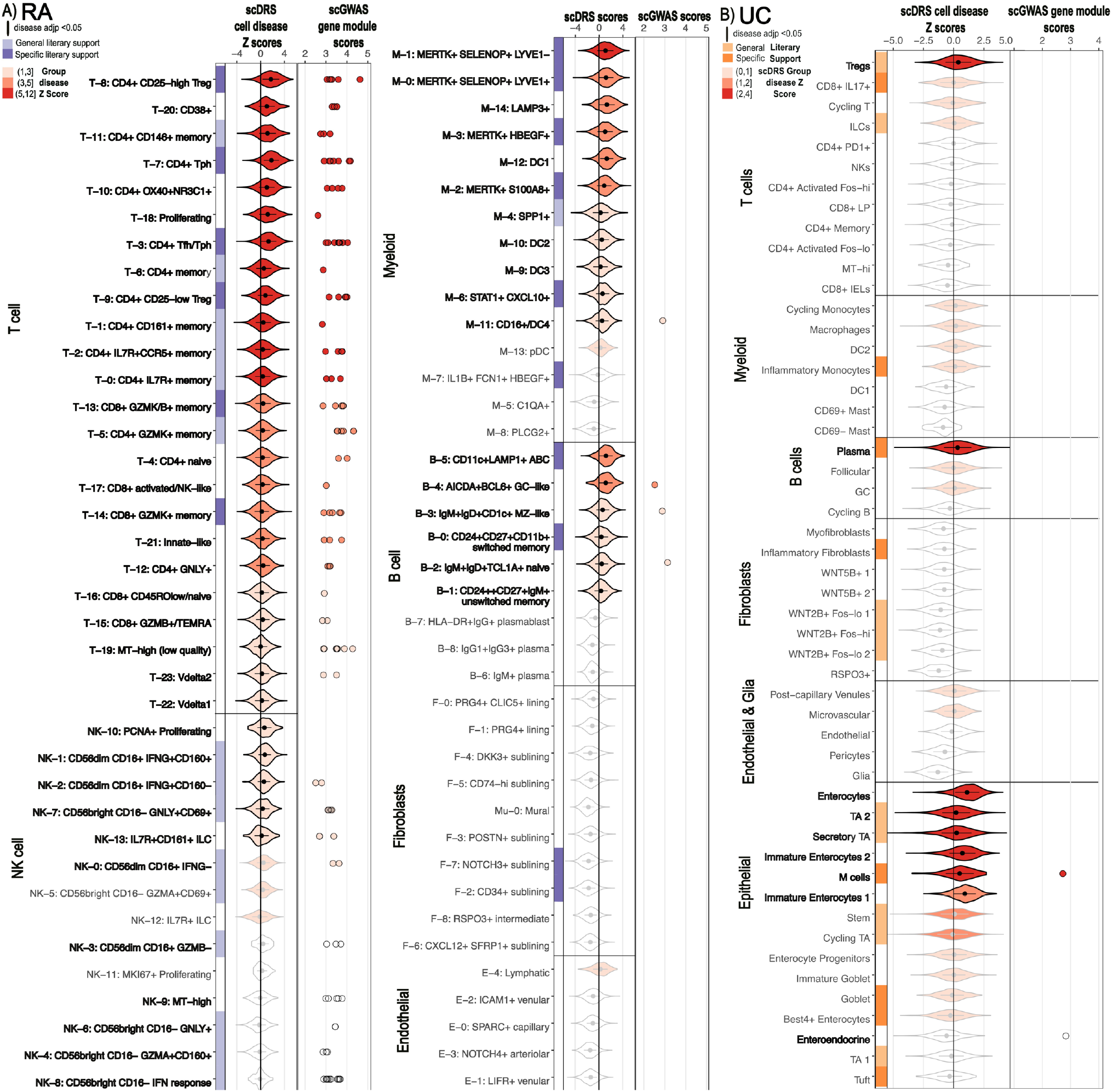
scDRS and scGWAS cluster significance results for RA and UC. For each cell-type and cluster, the single-cell level scDRS Z-scores are graphed with the violin plot colored and ordered according to the group scDRS Z-score (red scale). Nonsignificant cell clusters are outlined in gray while significant cell states are highlighted in black. scGWAS called gene modules and their disease scores are also plotted and are colored with the scDRS group Z-score gradient for easier comparison. Left: RA (rheumatoid arthritis). Right: UC (ulcerative colitis).

We next sought to evaluate the significant genes called by scGWAS and scDRS. Significant modules identified by scGWAS are clusters of genes that may represent a biological pathway and contain genes important for disease pathogenesis. scGWAS assessed these gene modules for each annotated cell type cluster. When performing gene set overlap analysis of the top ranked modules for RA, we found pathways strongly coordinating with the appropriate scRNA-seq cell state annotation(Mootha et al. 2003; Subramanian et al. 2005) (Supplemental Table 6). For example, T cell gene modules were frequently enriched with cytotoxic or T helper cell surface molecules while gene modules associated with NK cell states were enriched in genes involved in upregulating CD4 T cells and cellular responses to cytokines, chemokines, and cellular ligands. Many of these gene modules had overlapping genes and similar functions; despite having a total of 204 and 472 genes in NK and T cell cluster significant modules, there were only 63 and 87 unique genes, respectively. One gene in particular was found in nearly every significant gene module across cell states–CD2, which encodes for a surface antigen in all T cells and is involved with triggering T cells(“Immune Cell - CD2 - The Human Protein Atlas,” n.d.). Although scDRS does not provide cluster and disease based gene networks, it does identify genes whose expressions correlate with scDRS cell disease scores(M. J. Zhang et al. 2022). Gene set overlap analysis of the 100 genes most strongly correlated with scDRS disease scores across all cells showed enrichment of genes involved in differentiation and activation of T cells, general immune processes, macrophage-enriched metabolic networks, and plasmablastic cells. These findings reflect the high ranking of the corresponding cell states in scDRS disease scores and their strong population numbers. Importantly, to apply the same correlation analysis for a particular cluster, scDRS must be rerun with input specific to that cluster. When focusing on Tph cells (T-7), a phenotype identified in autoimmune diseases, highly correlated genes fit the role of helper T cells: Interleukin, cytokine, and TNFA signaling processes, and dendritic response to inflammatory stimuli processes(Pujana et al. 2007). In general, both MAGMA and module genes called significant by scGWAS have median positive correlation with scDRS disease scores across all cells (Supplemental Figure 2).

Furthermore, we explored if variance in significant genes might explain the different significant cell clusters identified by a gene-network based approach (scGWAS) and single-cell disease score approach (scDRS). First, we assessed if the reason that MERTK+ cells might not have been identified as significant by scGWAS is due to the tool’s reliance on predefined networks of genes. The MERTK cell relevant genes may not be connected in the network file used by scGWAS, so that MERTK+ cells are not called significant. Therefore, we evaluated if the genes that most highly correlated with scDRS disease scores for cells in the MERTK+ clusters were found in networks in the original scGWAS pathway file. Indeed, pairs of genes that are strongly associated with scDRS disease scores were connected, however, relationships between the genes beyond two were not supported and the 40 genes with the highest correlation to scDRS disease scores had only 6 pairings between them in the pathways file (Supplemental Table 7).

Similarly, we explored why the CD56 bright CD16-IFN response (NK-8) cells had the largest number of significant gene modules called by scGWAS while scDRS marked it with the lowest disease Z score across NK clusters and as not significant. As a controlled comparison, we also looked at a cell cluster with strong agreement between scGWAS and scDRS: CD4+ Tph (T-7). scDRS disease scores in all cells positively correlated with the expression of the NK-8 scGWAS module genes although T-7 scGWAS module genes had a slightly higher median correlation (0.08 vs 0.13) (Figure 3A,B). This relative increase was maintained when the eight genes identified by scGWAS as significant for both groups were removed (median correlations 0.005 NK-8 vs 0.02 T-7).

**Figure 3.**
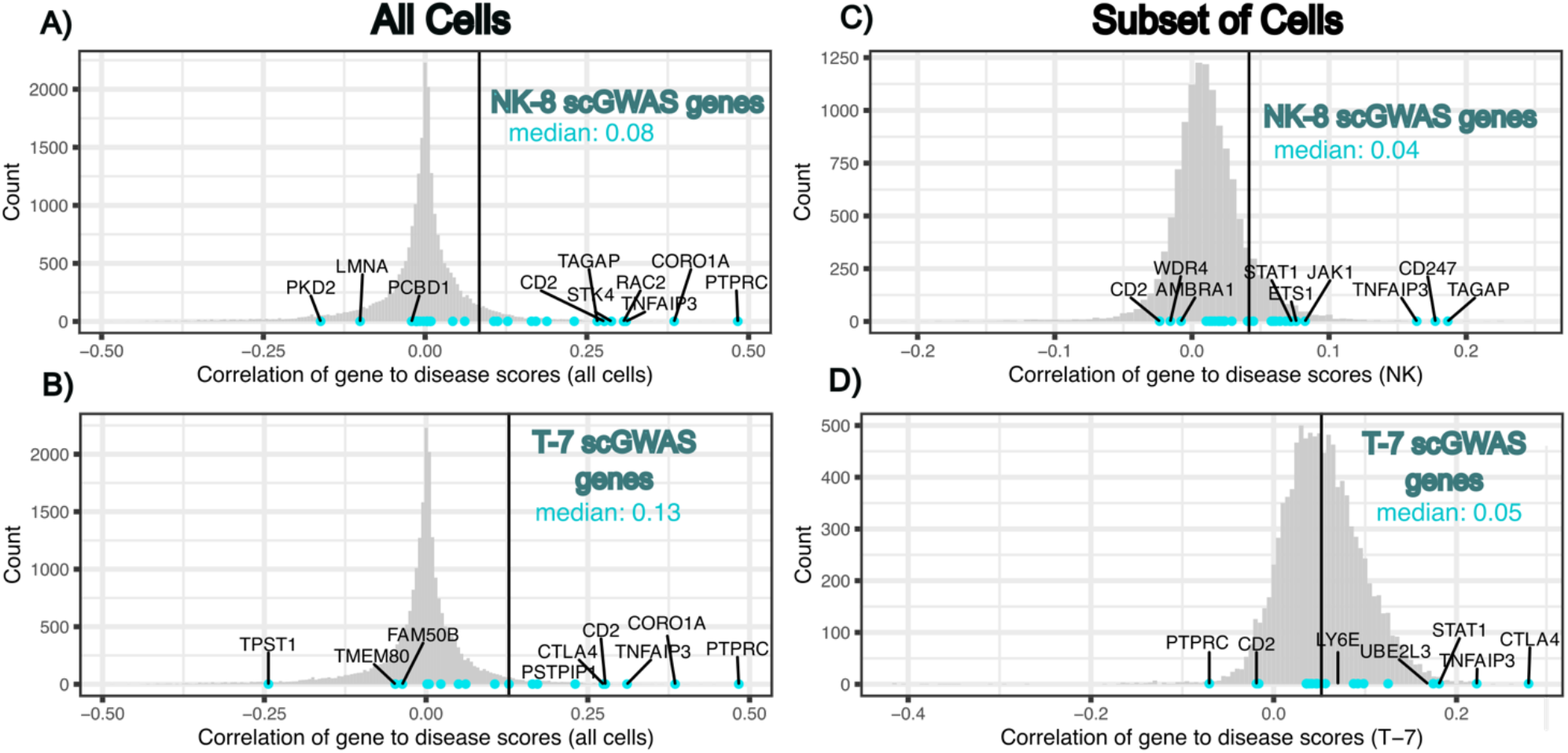
Histograms of the correlations of all studied genes with scDRS disease scores (grey) in all cells (A,B), NK cells (C), or T-7 cells (D) with the appropriately labeled scGWAS module genes highlighted (turquoise). Median correlation score of scGWAS genes is written and shown as a vertical line. scGWAS genes with highest and lowest scores are labeled.

Importantly, these correlations were comparable to that observed for all scGWAS genes as well as the top 100 genes ranked by MAGMA (medians of both 0.06) (Supplemental Figure 2B). The overall low correlations might be partly explained by using all cells rather than those specific to the cell state from which genes were identified. This explanation is supported by the low median correlation of MAGMA genes on which scDRS bases its analysis. However, median correlations decreased when only considering cells within the corresponding cell states (NK-cells & T-7) unlike those of the top ranking scDRS genes for each cell state (Figure 3C,D, Supplemental Figure 3). Still, these findings may be due to the high heterogeneity of disease scores among both of these cell groups, as defined by significant scDRS heterogeneity assessment, and the usage of of pseudobulk rather than single cell data by scGWAS. Historically, purely correlational approaches tend to be noisy and significantly impacted by the heterogeneity of the data (CITE). Therefore, to assess why CD56 bright CD16-IFN response (NK-8) but not other clusters had been called by scGWAS, we saw if the significant gene modules identified by scGWAS for this cell state show a higher expression in these cells compared to NK clusters without that same module called significant. The highest ranked cell state by scDRS, PCNA+ Proliferating (NK-10), had comparable expression of multiple NK-8 modules despite scGWAS not calling any significant gene modules for the cluster (Supplemental Figure 4). Instead, NK-8 had a higher expression of gene *STAT1* to many of the clusters, otherwise showing comparable patches of expression of the significant gene modules.

Although all scDRS additional features are outside the scope of this work, we evaluated the usage of the tools’s group-level metric to consider the heterogeneity of disease scores within a cluster(M. J. Zhang et al. 2022). This metric can hypothetically indicate if a provided cluster has inner-clusters of cells that should be further separated out based on the groupings of disease score. All large cell types in RA (T cell, B cell, Myeloid, NK, Fibroblast, Endothelial) had significant heterogeneous disease scores that positively correlated with the number of annotated clusters in each group (Supplemental Figure 5). Sixty seven of the seventy seven (87%) RA cell clusters had significant levels of heterogeneity in disease score, and the number of cells in a cluster had low positive correlation (adjusted R^2^ 0.29) with the heterogeneity Z-score and number of annotated clusters (adjusted R^2^ 0.37) (Figure 2, Supplemental Figure 5, **6, 7**). These findings suggest careful interpretation of a heterogeneity score in that statistically significant heterogeneity does not equal biologically significant heterogeneity. However, cell state annotations can be subjective and the non-significant p-values from our correlation analysis can be partly explained by low sample sizes. Despite these additional features and working at the single-cell level, scDRS was the most robust in memory usage and speed, although this is primarily due to the initial preprocessing step for scGWAS (**Table 2**). Notably, the number and size of cell states had a negligible effect on resource usage in both tools.

**Table 2.**
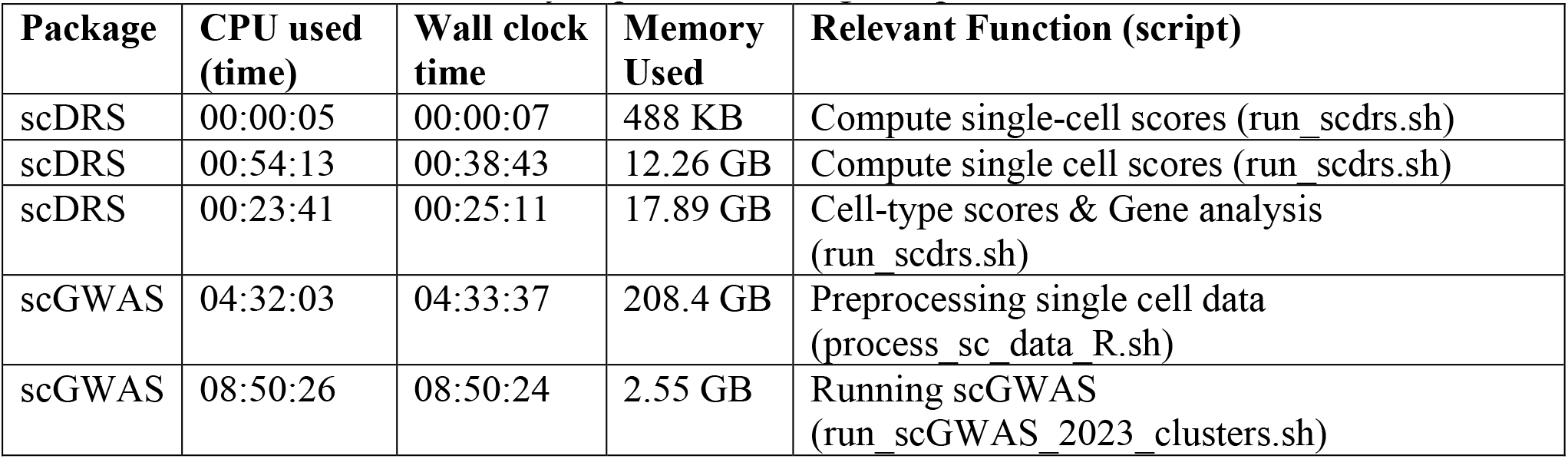
Resource usage of each package when running for the RA cluster-level data. Memory used refers to the max amount of memory required for a single step. Both were run with 15 CPUs.

### 3.2 scDRS allows successful distinguishment of similar diseases according to pathological cell clusters from the same scRNA-seq atlas

By using unique genetic information from diseases with the same scRNA-seq in relevant tissues, we demonstrate the feasibility to distinguish pathological cell clusters of similar diseases. We used summary statistics from GWAS for RAs and ankylosing spondylitis (AS) on the scRNA-seq data from inflamed RA synovial tissue to determine if scRNA-seq from a clinically similar disease can provide insight on disease-relevant clusters(F. Zhang et al. 2023; Ishigaki et al. 2022; Jiang et al. 2021). Similarly, we applied the GWAS statistics from UC and crohn’s disease (CD) on the scRNA-seq data from UC colon tissue(Smillie et al. 2019; de Lange et al. 2017). We found that while most T, myeloid, and B cell clusters were significant for RA, very few were significantly associated with AS (**Figure 4A**). Yet, cell states causally linked to AS according to a recent Mendelian randomization study were all called significant: CD8+ activated/NK-like (T-17), pDC (M-13), and unswitched memory cells (B-1)(Fei et al. 2024). AS and RA showed the greatest differences across the T, NK, and myeloid cells. While essentially all T cell clusters showed significance for RA, only CD8+ activated NK-like (T-17) and proliferating (T-18) T-cells showed significance for AS, following literature supporting the role of mostly CD8+ T cells in AS(Gracey et al. 2020; Xu et al. 2021). Conversely, far more NK cell clusters were called significant for AS with again support from literature. GZMB was at much lower levels in AS patients in previous NK-focused scRNA-seq analysis and ELISA studies, supporting the significance of CD56dim CD16+ GZMB-cells (NK-3)(Ren et al. 2022). Similarly, the significantly called IL7R+ ILC (NK-12) cluster showed similar upregulation of genes, including IL7R, as a NK cluster upregulated in AS according to previous single cell analyses(Ren et al. 2022; Liu et al. 2021). Finally, most of the CD56bright CD16-(NK4,6,8) NK cell clusters were called significant, which is supported by the upregulation of CD56bright NK cells in AS(Ren et al. 2022; Liu et al. 2021). These differences were consistent when using different MAGMA windows (Supplemental Figure 8). Although fewer clusters were called significant for UC and CD (8 and 6, respectively) (Figure 4B), we still saw relevant differences in pathological cell types. These findings contrast the few calls from original scDRS analysis with the same summary statistics when using larger scale cell groups(M. J. Zhang et al. 2022). Epithelial cells linked to UC and fibroblasts linked to CD most clearly distinguish the diseases, a finding maintained when using different MAGMA windows (Supplemental Figure 9). For example, we found that NK cells, CD4+ activated, and CD8+ lamina propria (LP) cells were enriched in CD compared to UC while only Tregs, CD8+ IL17+, and Cycling T cells were enriched in UC. Indeed, the enrichment of specifically CD8+ LP cells, NK cells, and activated CD4+T cells has been supported by independent CD single cell analyses(Jaeger et al. 2021). Importantly, we also captured the unique genetic support of fibroblast enrichment for CD, first with calling RSPO3+ fibroblasts significant when multiple CD specific SNPs have previously connected this phenotype(Jostins et al. 2012). Similarly, our findings for WNT2B+ fibroblasts match the fact that the group has only shown genetic connection to CD despite it being expanded in both diseases(Burke et al. 2007; D’Alessio et al. 2022).

**Figure 4.**
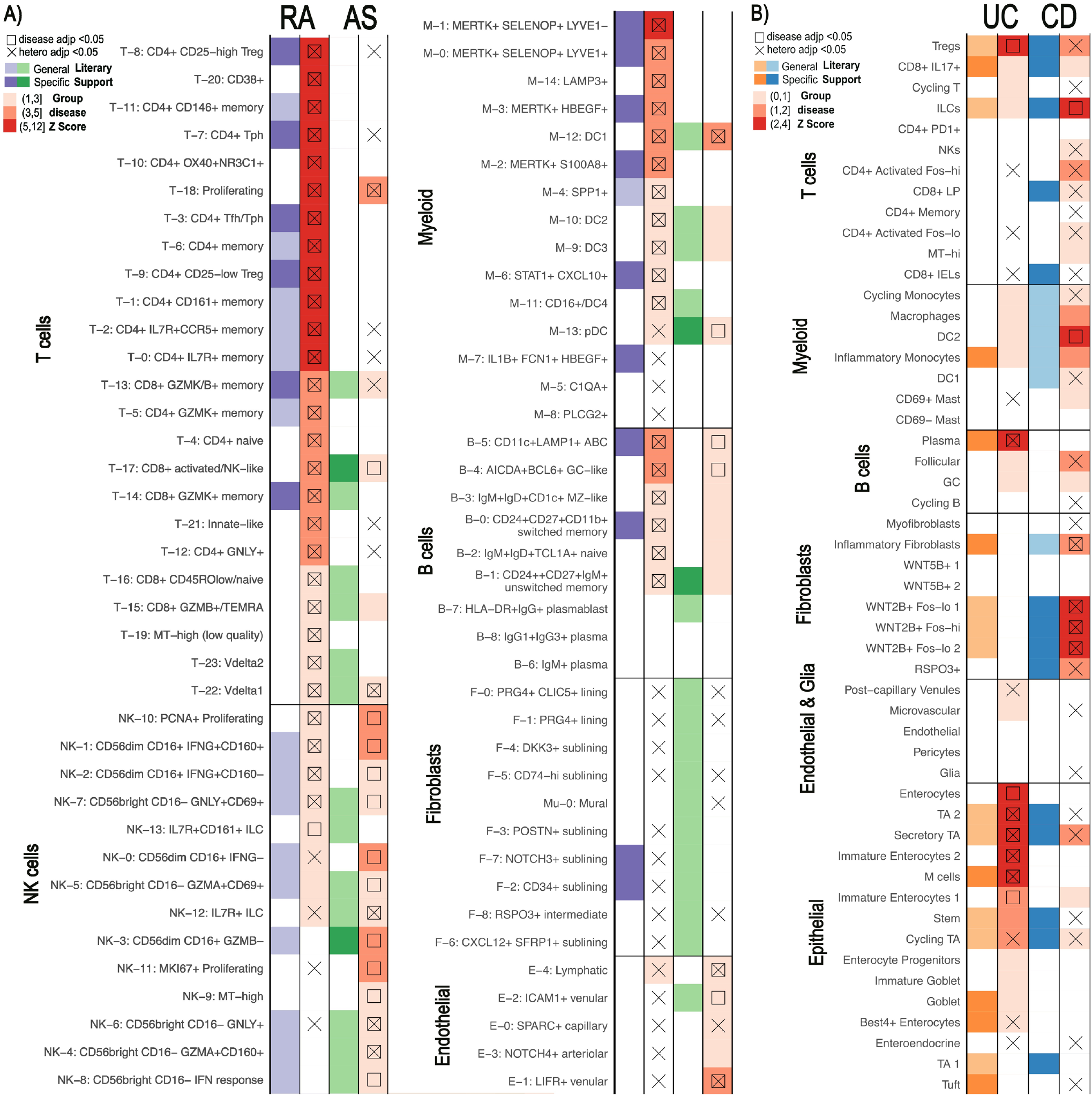
Comparison of similar diseases with scDRS. Summary statistics unique to each disease were used on the same scRNA-seq data for each pair (F. Zhang et al. 2023; Smillie et al. 2019). scDRS defines significant clusters (annotated according to original papers) with a group disease Z-score as shown in the gradient legend. Cell clusters with literary support for either disease are labeled in purple/orange for RA/UC and green/blue for AS/CD, respectively. General literary support means that a cell type with multiple cell states is supported by the literature while specific means a specific single cell state was supported. Left: Rheumatoid arthritis (RA) vs Ankylosing Spondylitis (AS). Right: Ulcerative Colitis (UC) vs Crohn’s Disease (CD).

### 3.3 Positional SNP-Gene linking methods provide greater statistical power than the tested alternatives

Methods integrating scRNA-seq and GWAS summary statistics rely largely on the same preprocessing steps, yet a standardized guidance for these steps is lacking. Therefore, we evaluated the impact of inputs and preprocessing steps on results identified by scDRS due to its high sensitivity.

First, we considered the robustness of results when using solely positional information to connect noncoding SNPs to genes. The primary positional method to link SNPs to genes is MAGMA which relies on a window size parameter determining the distance a SNP can be from a gene to be incorporated(Leeuw et al. 2015). Because there is no standardization on MAGMA window size beyond the notion that a larger window size incorporates SNPs falling in cis-regulatory elements, we evaluated the impact of the most used window sizes on results (details in Methods) (M. J. Zhang et al. 2022; Jia et al. 2022; Watanabe et al. 2017; Sobrin et al. 2022; Hariharan and Dupuis 2021; Watanabe et al. 2019; Skene et al. 2018; Zhu and Stephens 2018; Finucane et al. 2018). Different window sizes for RA analyses only changed the significance calls for 16 of the 77 cell clusters in at least one of the window-sizes, half of which are only different in one window size (Figure 5). Importantly, none of these clusters had the top 20 group disease scores in our original results (50-35kb window). There also did not appear to be a clear pattern across the window sizes in terms of the numbers of significant clusters or the clusters changed in significance. These findings were similar with diseases beyond RA, with results for CD having the greatest differences across window sizes (Supplementary Figures 6, 7). This robustness might be explained by the finding that although only 54% of genes were shared across the top 1000 MAGMA ranked genes in all window sizes, these shared genes consistently had most of the lowest p-values (Supplemental Figure 10).

**Figure 5.**
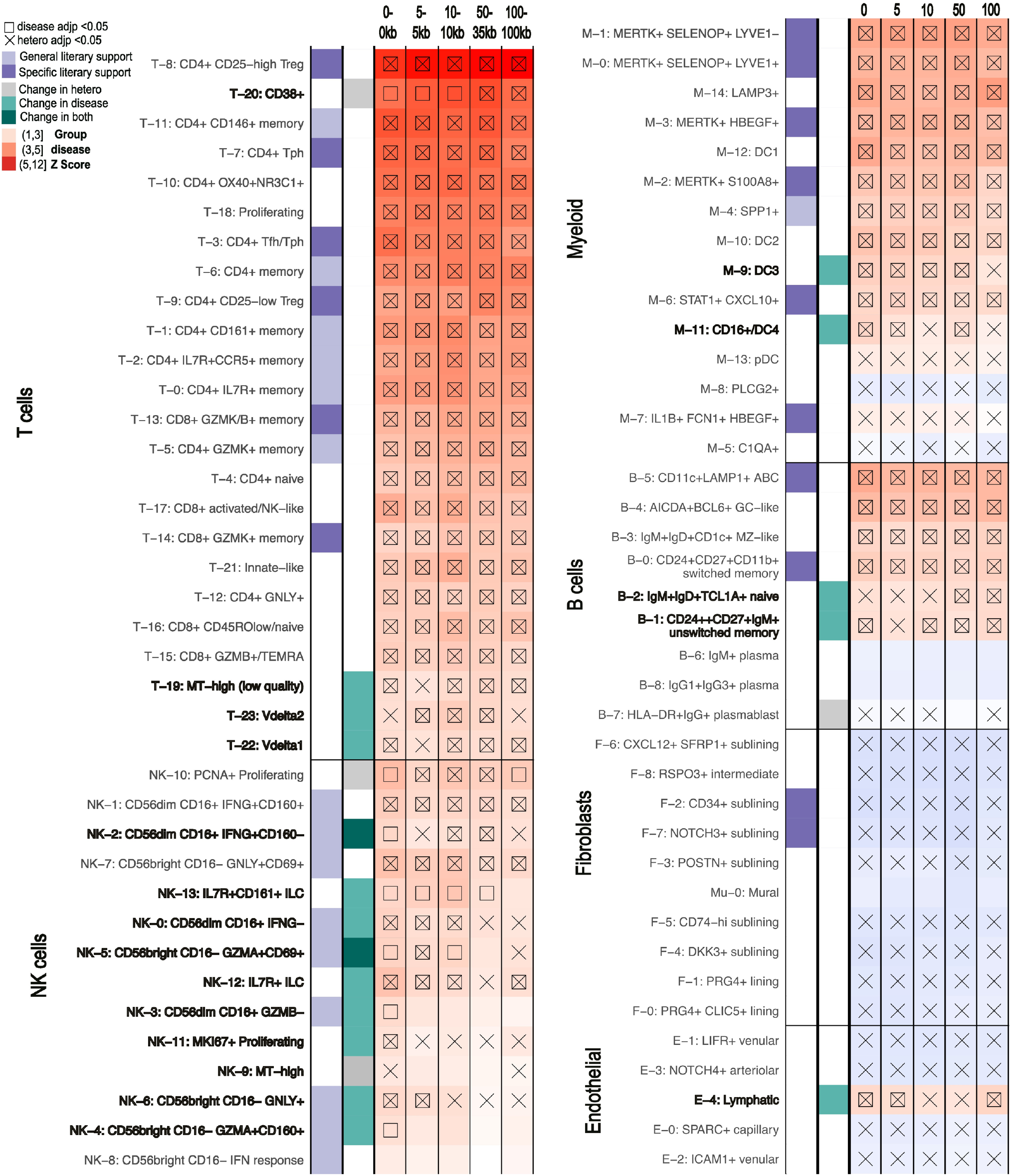
scDRS results for RA of clusters that show different levels of significance with different MAGMA windows being used to generate the GWAS inputs (0-0kb, 5-5kb, 10-10kb, 50-35kb, 100-100kb). scDRS defines significant clusters with a group disease Z-score as shown in the gradient legend (significant scores marked with square). Cell states with significant heterogeneity scores are marked by an X. General literary support means that a cell type with multiple cell states is supported by the literature while specific means a specific single cell state was supported. Cell states with changes in just scDRS disease score, heterogeneity score, or both significance calls across MAGMA windows are marked in bold and with grey or turquoise squares.

Given the growing concern over positional methods inaccurately assigning SNPs to genes, we next explore the usage of non-positional based data within the framework of FUMA. Although tools can be found in Table 3, we focused on FUMA as the commonly used tool that incorporates eQTL and chromatin contact data along with positional information from MAGMA to express summary statistics at the gene-level(Leeuw et al. 2015; Watanabe et al. 2017; Gazal et al. 2022; Weeks et al. 2023; Yang, Chen, and Zhao 2021). While it uses a different summary statistics processing that doesn’t allow direct comparison to our own MAGMA based analyses, we were still able to use its MAGMA to consider the impact of alternative linkage methods (details in Methods). The 1000 genes with the highest significance calls were significantly different between methods, even when comparing MAGMA run on FUMA summary statistics vs our summary statistics (Supplemental Figure 11). When just considering genes supported without MAGMA, only 445 genes were output as significant (Supplemental Table 8). This relatively small number follows the trend of non-positional based methods providing significantly smaller genesets with their focus on highly confident linkages due to noisy data sources (Table 3). Importantly, the authors of scDRS found that using less than 1000 genes causes a significant reduction in power despite it being one of the most sensitive methods available given its analysis at the level of single cells(M. J. Zhang et al. 2022). Indeed, FUMA analysis that combined positional with non-positional methods showed similar results to purely using MAGMA but with only 28 of the 52 original clusters called significant (Supplemental Figure 12). This method also added 2 significant clusters: HLA-DR+IgG+ plasmablasts (B-7) and MKI67+ Proliferating NK cells (NK-11), which were still not called significant when increasing the MAGMA window size to 100kb, a size commonly used to capture cis-regulatory element SNPs (Figure 4).

**Table 3.** Current methods to link SNPs to genes and the estimated number of genes output, form of significance output, and interface. All tools address linkage disequilibrium.

| **Name (Year)** | **Method** | **Est. Gene list size** | **Score** | **Interface** |
| --- | --- | --- | --- | --- |
| **cS2G** (2022) | Linear combination of linking scores from main S2G strategies, exon, promoter, eQTLGen, and GTEx cis-eQTL, EpiMap, ABC, and Cicero. Restricts each strategy to gene w/ highest linking score. | <500 (depends on # lead variants) | cS2G score | Scripts provided |
| **PoPs** (2020/23) | Similarity based filtering of MAGMA results (although paper described other input options). | <200 (depends on # lead variants) | PoPs score (for relative ranking) | CLI |
| **nMAGMA** (2021) | Network-enhanced MAGMA links SNPs to genes by considering tissue specificity (Hi-C and eQTL) and functional interactions (WGNCA), then use MAGMA to get significance of genes. | 1000+ | Z-scores and P-values | Scripts provided |
| **FUMA** (2017) | SNP2GENE Module: Identifies lead SNPs, can run MAGMA or map using eQTL, position, and chromatin-interaction | MAGMA based 1000+, otherwise <700 | MAGMA Z-scores/P-values or min p-value of linked SNPs | Web tool |
| **MAGMA** (2015) | Maps SNPs to genes via positional window, empirical gene p-value via permutation followed by PCA regression | 1000+ | Z-scores and P-values | CLI |

## 4 Discussion

In this study, we evaluated two representative computational strategies for linking genetics to single-cell phenotypes according to the enrichment of literature supported calls, robustness, and interpretability of results. Although both strategies identified verifiable significant cell states, the single-cell based scDRS identified the greatest number with literary support. Indeed, our findings helped newly confirm genetic-phenotype predictions previously made according to purely transcriptomics or genetic analyses in both RA and CD, specifically. Given the increased power, efficiency, and exploratory analysis provided by single-cell methods, we sought to see if bulk-based gene network methods benefit gene-centered analyses. Indeed, scGWAS is more distinctly built to identify probable gene sets relevant to pathological cell states while scDRS is targeted more towards single cell based exploration(M. J. Zhang et al. 2022). scDRS provides genes correlating with the disease scores of all cells provided, requiring repeated runs to get the correlating genes for each cell type. Results are then largely affected by the heterogeneity of the dataset, with correlation ranks being biased by proportions of cell phenotypes within the dataset. However, scGWAS is also confined by the provided network file. While removing false positives by requiring a known set of connected genes to have increased expression compared to single genes, the algorithm also assumes that the pathway file contains all possibly relevant gene connections. Therefore, true positives can be lost such as with the MERTK+ cells and the reasoning behind selective calls can be unclear as in the case of NK cell states for RA. Additionally, many of the significantly called scGWAS gene modules overlapped, depleting information content, perhaps due to the lack of cell type specificity of the pathways. This finding further clarifies the importance of not necessarily using the number of significant gene modules identified as a relative metric of significance for a cell type(Jia et al. 2022). Although scGWAS provides gene modules more conducive for certain analyses, the original network file should be largely considered according to a researcher’s specific focuses.

In our work, we also successfully identified fine-scaled significant clusters for diseases without their scRNA-seq atlases by applying these tools to atlases for similar diseases. Importantly, the results still allowed clear distinction between the two diseases, with literary support for the found differences from other single-cell based analyses. Publicly available scRNA-seq data is not always available or sufficient for a certain disease, so instead researchers might need to apply the existing and relevant GWAS summary statistics to the scRNA-seq data generated from a clinically similar disease. Our findings support the ability for researchers previously constrained by the lack of appropriate scRNA-seq atlases to study diseases while not sacrificing fine-scale analyses.

Finally, we also evaluated methods incorporating noncoding SNPs for identifying pathogenic cell states. Although we found that MAGMA based results arse robust to window sizes, we also considered non-positional methods to link SNPs to genes with FUMA and found the decreased power from these tools have significant impacts on results. Ideally, one would be able to combine strict window MAGMA results with that of a non-positional method, however the need to combine different significance scales complicates this. The p-values output by FUMA and similar methods also often do not account for the uncertainty in the predicted SNP-gene linkages as done by MAGMA. For now, if using tools reliant on a long list of genes, we suggest focusing on cell types consistent across window sizes for MAGMA and manually considering genes called by other tools like FUMA. It’s important to note that regardless of the window sizes used, many SNPs were still not assigned to a with MAGMA gene. For example, with a moderately large window size of 50-35kb, about 60% of SNPs for RA and UC were linked to a gene which decreased to about 40% when that window was reduced to 10-10kb. Outside of these methods, repeating analyses with multiple GWAS summary statistics and scRNA-seq cohorts is equally relevant to ensure repeatability of results.

One way to circumvent linking SNPs to genes is using cis-regulatory elements (cREs) SNPs fall in directly. Given cRE activity is highly dependent on cellular behavior and allows more accurate deconvolution of cell types, this switch could also allow separation of more nuanced cellular states(Maas, Sigauke, and Dowell 2023). Additionally, tools like Cicero link cREs to their regulated genes from single cell data(Pliner et al. 2018). While classic scRNA-seq data cannot usually capture the activity of these elements, scATAC-seq and 5’ scRNA-seq have provided valuable alternatives. scATAC-seq has been used to predict relevant cell states by linking SNPs to open chromatin regions, but no standardized pipeline has been developed(Pliner et al. 2018; Scherer et al. 2021; Orchard et al. 2021). This is likely due to the higher cost of scATAC-seq compared to scRNA-seq and the disconnection between open chromatin and regulatory activity(Corchete et al. 2020). Conversely, Moody et al. successfully applied 5’ sc-RNA-seq to detect the transcription of cREs and genes simultaneously and developed a metric to identify cell types enriched in trait heritability(Moody et al. 2023). Interestingly, they used the same summary statistics as our work for crohn’s disease (CD) and ulcerative colitis (UC). Despite using gene based methods, we were still able to capture the fibroblast and dendritic cell enrichment for CD that they found. However, unlike their results, we did not find an overall enrichment of T/NK cells in UC compared to CD but found some specific states in these cell types oppositely enriched and supported by the literature(Jaeger et al. 2021). These differences can be explained by the fact that Moody et al relied on general lymphocyte 5’-scRNA-seq for analysis while we used scRNA-seq specifically from the colon mucosa of UC patients. The cell states we identified as seeming to conflict with larger scale findings from Moody et al are unique to intraepithelial lymphocytes and likely would not have been found in their data. Overall, these results showcase the need for careful interpretation when relying on non-disease tissue specific scRNA-seq data; this warning would be particularly applicable to cRE based methods given how specific regulatory element activation is to cell-states. Exciting insight will come from evaluating the adaptation of algorithms like scDRS and scGWAS to the growing cRE-based single cell data.

Overall, although 5’-scRNA-seq allows improved detection of cREs, these methods still cannot capture most cREs compared to tools like nascent-RNA-sequencing which are still under progress to being applied at the single cell level (Moody et al. 2023; Whalen, Truty, and Pollard 2016; Lizio et al. 2015; Sigauke et al. 2023). A clear theme from our analyses was that the usage of fine-scale scRNA-seq atlases allowed distinctions between nuanced cell states supported by genetic causality more biologically informative compared to the general atlases used previously. Indeed, both scGWAS and scDRS struggled to note the significance of large cell groups if only a small portion of the cell type, such as a nuanced cell state, is significant. Using tissues and nuanced annotations specifically relevant to the disease can help circumvent the decreased sensitivity with heterogeneity. The heterogeneity score from scDRS might also help suggest when more nuanced cell states are required. While disease-specific and fine-scaled single-cell cRE atlases continue being developed, tools like scGWAS and scDRS provide key opportunities for taking advantage of the vast set of sc-RNA-seq atlases available. Indeed, we’ve found that these tools can even allow single-cell level analyses for diseases without fine-scaled sc-RNA-seq atlases currently accessible if an atlas is available for a similar disease. The development of tools like scDRS, scGWAS, as well as improved SNP-Gene-cellstate linking methods provide key steps for identifying biological targets for treatment development.

## Supporting information

Supplemental_Mat_Figs

## 6 Author Contributions

HT and FZ brainstormed the original project and HT devised the final focuses and design. KR ran scGWAS on the relevant data. HT did all other analyses. HT took the lead in writing the manuscript with support from KR. JI advised on GWAS related analysis and integration. FZ supervised the project with the aid of JI. LV provided feedback in the final manuscript editing. All authors provided critical feedback and helped shape the research, analysis and manuscript.

## 7 Acknowledgments

This work was supported in part by the Interdisciplinary Quantitative Biology (IQ Biology) PhD program at the BioFrontiers Institute, University of Colorado Boulder, and the National Science Foundation NRT Integrated Data Science Fellowship (award 2022138). The PhRMA grant and the Arthritis National Research Foundation grant to F.Z. The Curci Scholarship from the Shurl and Kay Curci Foundation (to H.T.) also enabled this work. We appreciated the constructive feedback from the Zhang Lab members. This work would also not have been possible without the IT support from CU-Anschutz Medical Campus.

## 8 Data Availability Statement

The GWAS summary statistics used in this study can be found in supplementary table. The single-cell count data for RA is available on Synapse (https://doi.org/10.7303/syn52297840). The single-cell count data for UC is available at the Single Cell Portal (SCP259). The Code used for analysis are available at [https://github.com/fanzhanglab/SCRNA-GWAS-Benchmarking].

