## Supplemental_Mat_Figs for "Single-cell based integrative analysis of transcriptomics and genetics reveals robust associations and complexities for inflammatory diseases"

Supplementary Material

### 1 Supplementary Notes

**Methods not considered in analysis:** CocoNet uses a similar network-based approach to the more recent scGWAS that boasts more statistical robustness. Similarly, sc-linker was published by the same laboratory as that which produced scDRS but does not perform at the single-cell level. Sc-linker instead focuses on the usage of enhancer-gene linkage methods to link SNPs to genes rather than classic positional methods; a consideration already addressed in this study.

**Gene universes:** Regardless of the tool used to map SNPs to genes, the genes must be used in both the scRNA-seq data and mapping tool. Genes identified as relevant by MAGMA or other methods may not have been considered in the scRNA-seq and therefore not be incorporated in analysis. Although only 1.2% of the MAGMA genes used for scDRS with RA had no known aliases identified in scRNA-seq genes, 18% of the top 1000 genes identified by MAGMA for UC had no known aliases found in the scRNA-seq data (Supplemental Table 5). For example, the scRNA-seq from UC did not include many genes in the 1000 top ranked by MAGMA. This list included the Mast cell growth factor *IL3*, and fibroblast growth factor *FGF21*. Both of the cell groups linked to these factors were not called significant by scDRS, possibly due to an inability to consider these genes in the analysis rather than their true lack of significance. Although remapping sequences to genomes including the annotated genes is possible, this introduces analytical differences that may disrupt the published cell state annotations and results.

### 2 Supplementary Figures and Tables

Supplementary Tables can be found in a single excel file labeled Supplementary_Material.xlsx where each sheet corresponds to the following:

Supplemental Table 1: Literature support for cell states in rheumatoid arthritis and ankolyzing spondylitis

Supplemental Table 2: Literature support for cell states in ulcerative colitis and crohn’s disease

Supplemental Table 3: GWAS summary statistics sources and metadata

Supplemental Table 4: Windows used found for positional SNP-gene linking tools from literature search

Supplemental Table 5: Differences in genes available and linking for RA and UC between GWAS (based on MAGMA) and scRNA-seq before and after addressing gene aliases.

Supplemental Table 6: Significant gene modules from scGWAS, module Z score, and notes from GSEA.

Supplemental Table 7: The 30 genes whose expression have the highest correlation with disease scores of MERTK+ cells and the gene to which they can be paired by the original scGWAS pathway network file with the highest correlation.

Supplemental Table 8: P-value cutoffs of the genes output by either FUMA or MAGMA

#### 2.1 Supplementary Figures

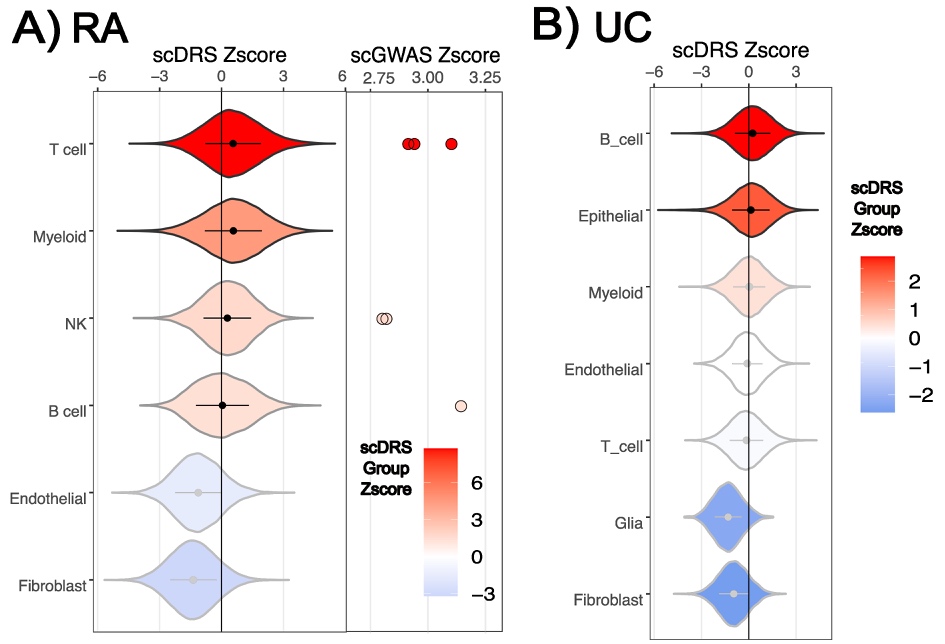

#### Supplemental Figure 1. Large-scale cell type results for RA and UC. For each cell-type, the single-cell level scDRS Z-scores are graphed with the violin plot colored and ordered according to the group scDRS Z-score. Nonsignificant (p>0.05) cell clusters are outlined in gray. Cell types significant only with p(<0.1) have the mean summary plotted in black (NK & B-cell for RA, Myeloid for UC). scGWAS does not directly assign cell-type significance scores but instead scores gene modules found in the clusters. The significant gene-module scores are plotted with the color gradient still following the scDRS group Z-score for easier comparison (UC had no significant modules).

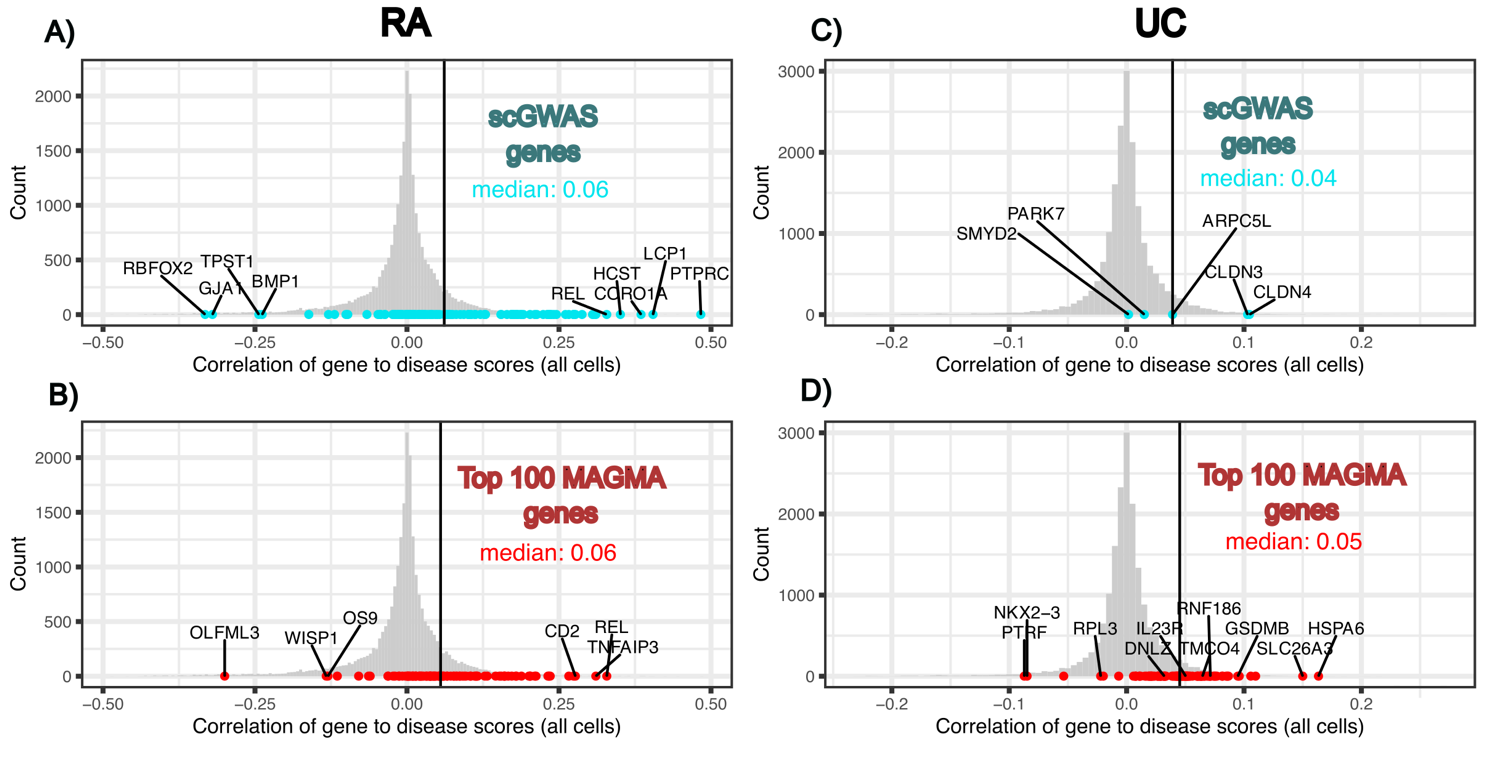
**Supplementary Figure 2.** Histograms of the scDRS correlation scores of all studied genes with disease scores in all cells with the scGWAS module genes or top 100 MAGMA genes highlighted by color (scGWAS=turquoise, MAGMA=red). The scGWAS or MAGMA genes with the highest and lowest scDRS correlations, and the scGWAS & MAGMA genes are annotated. The median correlation of the highlighted genes is listed. RA=rheumatoid arthritis, UC=ulcerative colitis.

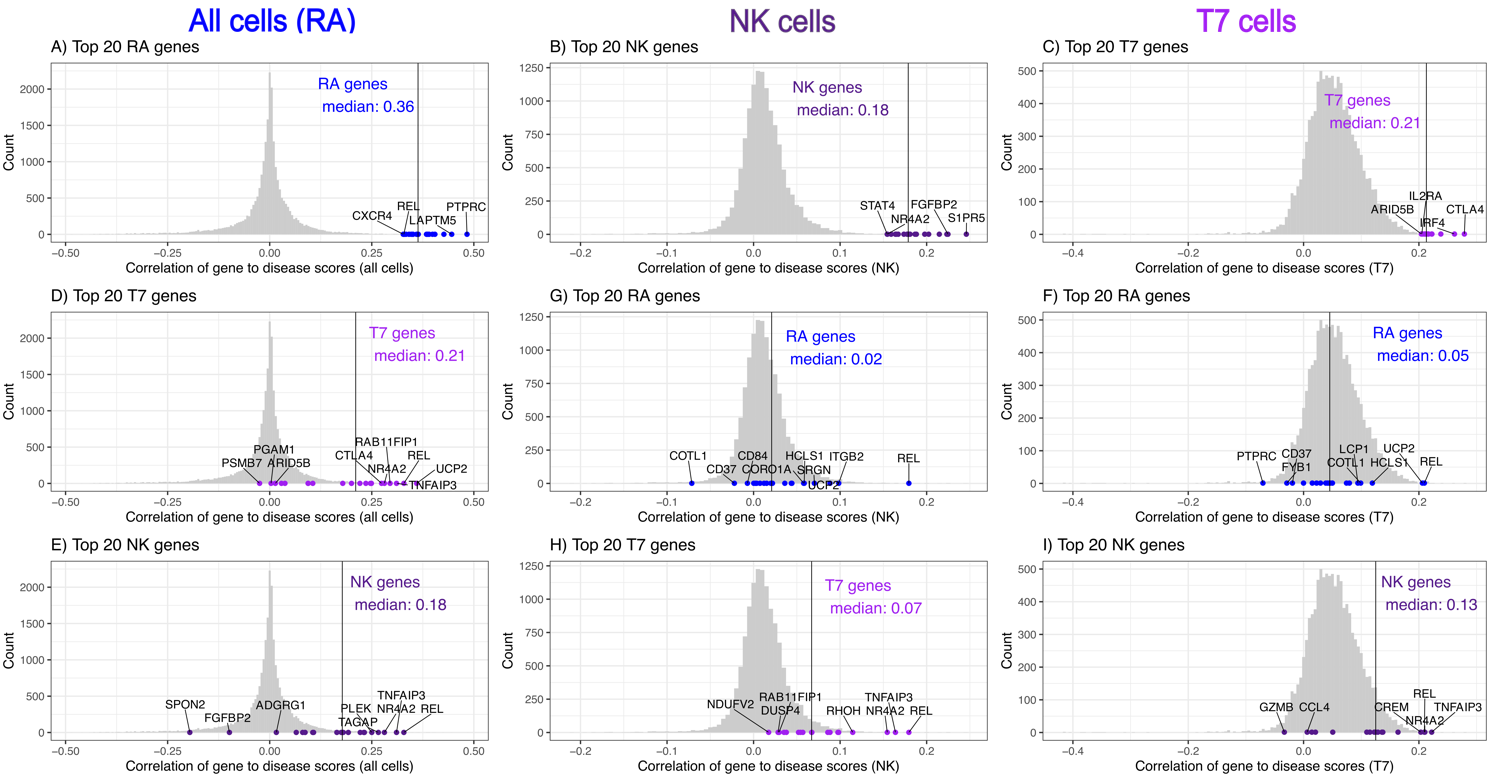
**Supplementary Figure 3.** Histograms of the scDRS correlation scores of the top 20 genes correlated with disease scores in all cells (RA), NK cells, or T-7 cells, within the different cell type options. The genes with the highest and lowest scDRS correlations within the top 20 list are annotated. The median correlation of the top 20 genes are listed. All data is from the RA analysis.

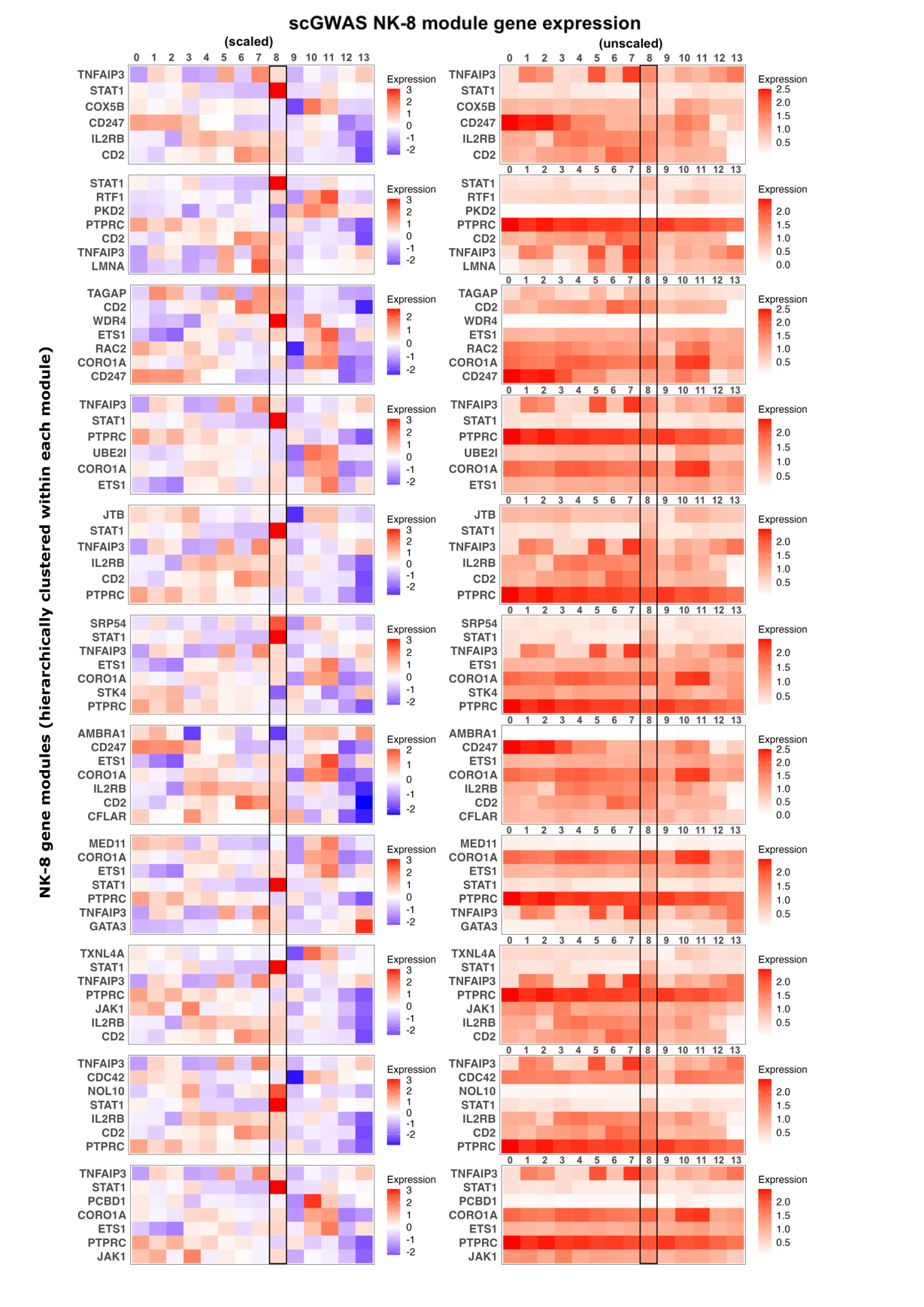

**Supplemental Figure 4. Heatmaps of expression of NK scGWAS significant modules genes.** Mean expression of the 11 NK-8 significant module genes according to scGWAS in both scaled expression and raw mean expression (counts) for the 14 NK clusters.

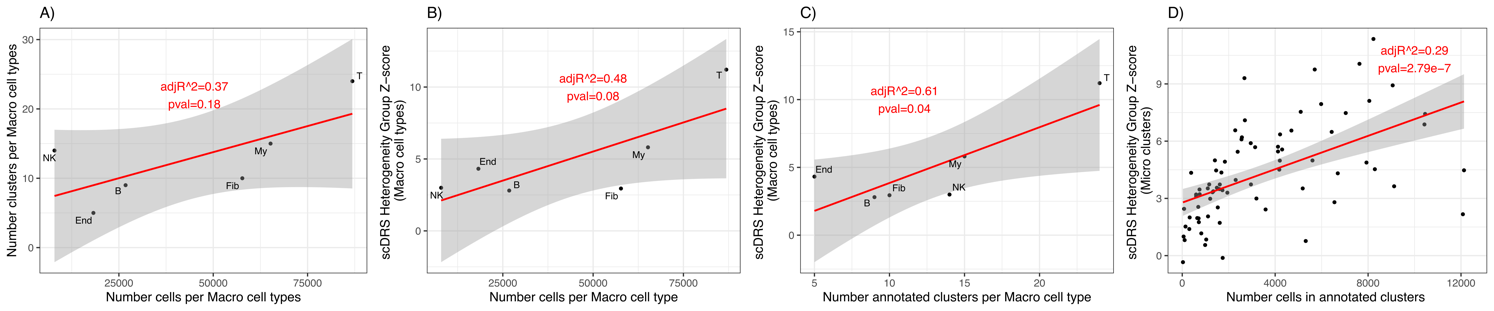
**Supplemental Figure 5.** Linear regression between heterogeneity score and number of clusters/cells in RA scRNA-seq data. The adjusted R^2^ and model p-value (F-test) are included. Left: the number of cells in Macro-cell types (T-cell, B-cell, Myeloid (My), NK, Fibroblast (Fib), Endothelial (End))  and their number of annotated clusters (A) and group scDRS disease score heterogeneity z-scores (B) (N=6). C) Number of annotated clusters in large-cell types and group scDRS disease score heterogeneity z-scores (N=6). B) the number of clusters in large-cell types and their group scDRS disease score heterogeneity z-scores. D) Number of cells in annotated clusters, and their group heterogeneity z-scores (N=77). Details of the linear regression results can be found in the jupyter notebook heterogeneity.ipynb on github.

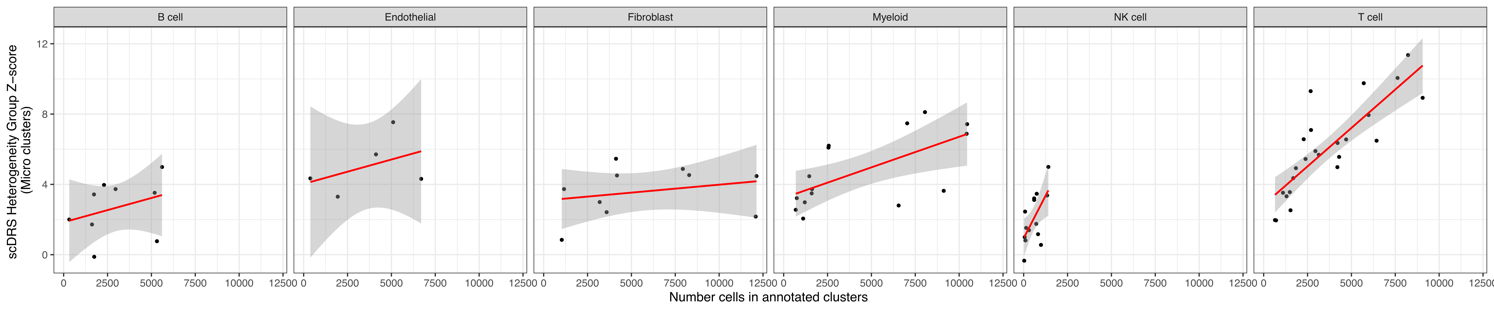
**Supplemental Figure 6.** Linear regression between heterogeneity score and number of cells in each cluster annotated from RA scRNA-seq data, separated by cell type. The adjusted R^2^ and model p-value (F-test) are included. Details of the linear regression results can be found in the jupyter notebook heterogeneity.ipynb on github.

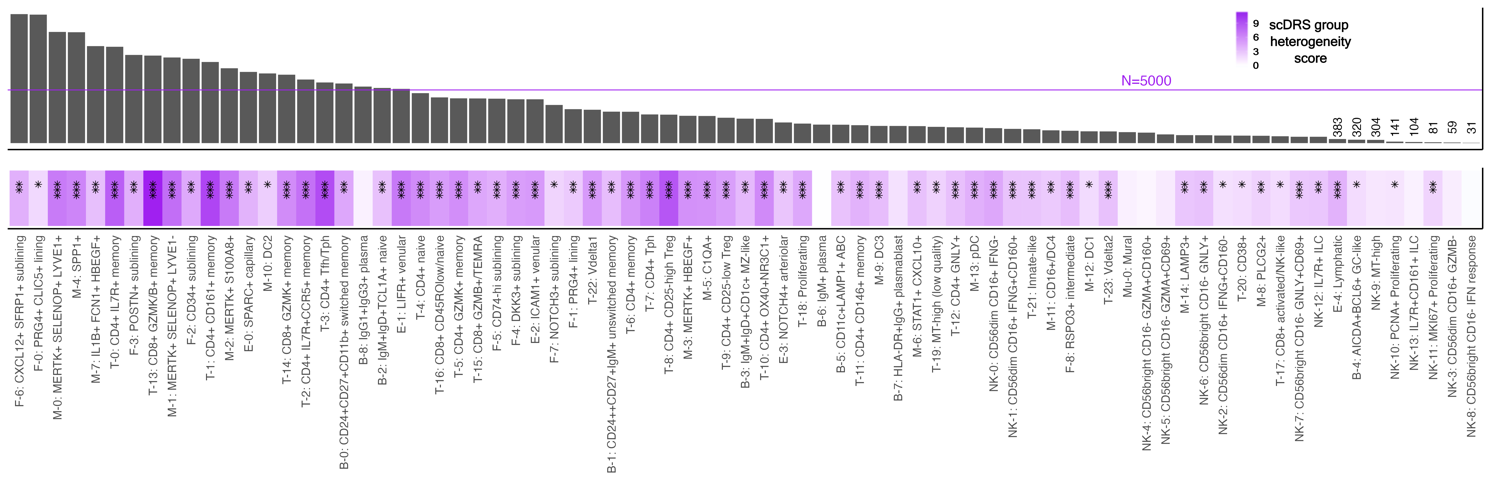
**Supplemental figure 7:** The scDRS group heterogeneity scores [of disease scores] of cell clusters from RA and the size of said clusters. Any cluster below 500 cells is noted by the number of cells while the rest have a N=5000 bar for reference with the largest number being 12k. Significance legend: *P<0.05, **P<0.01, ***P<0.001.

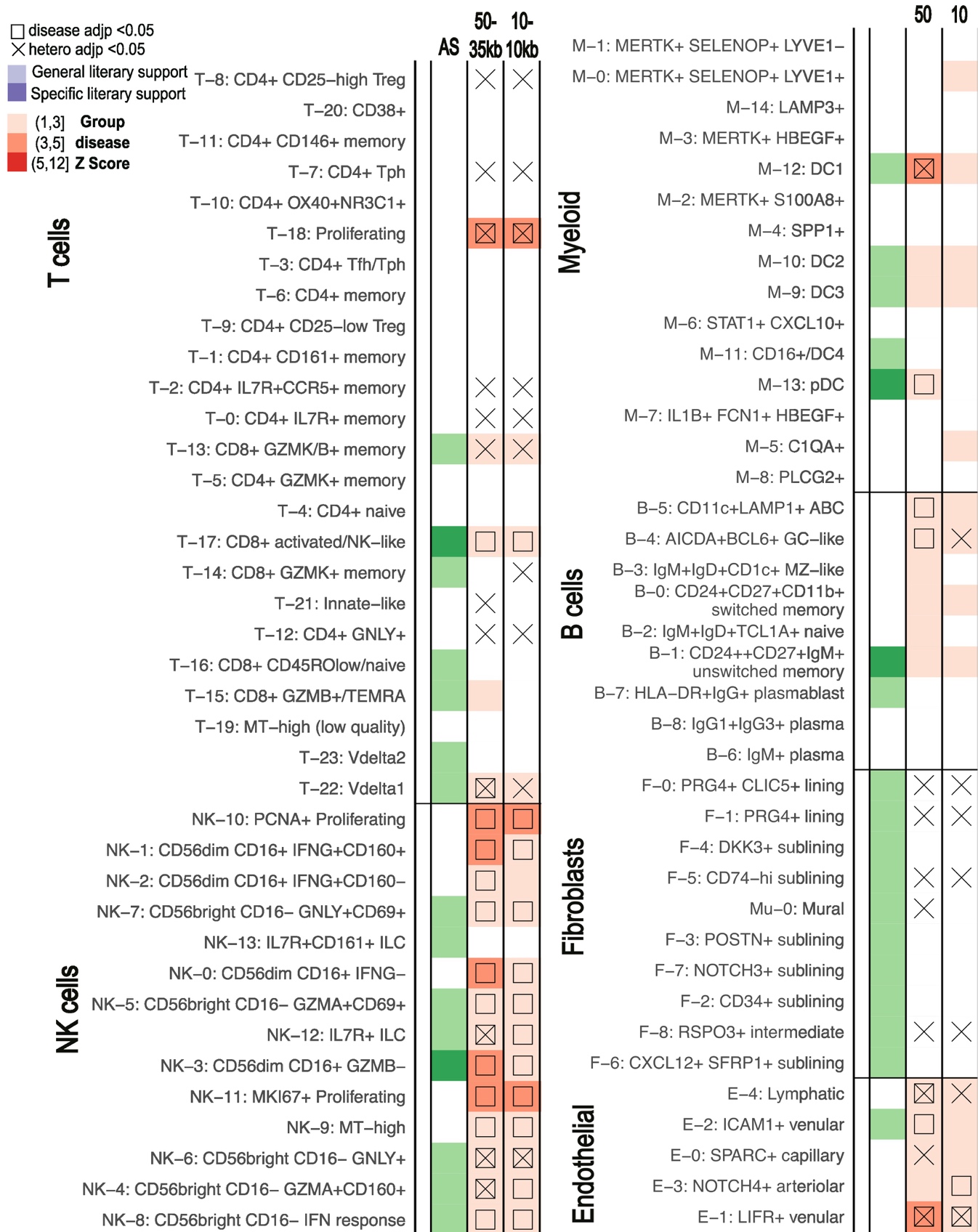

**Supplemental Figure 8. MAGMA window comparisons for ankylosing spondylitis.** scDRS results of significant clusters for AS using 50-35kb and 10-10kb windows. scDRS defines significant clusters with a group disease Z-score as shown in the gradient legend. Significance legend: *P<0.05, **P<0.01, ***P<0.001. AS = ankylosing spondylitis.

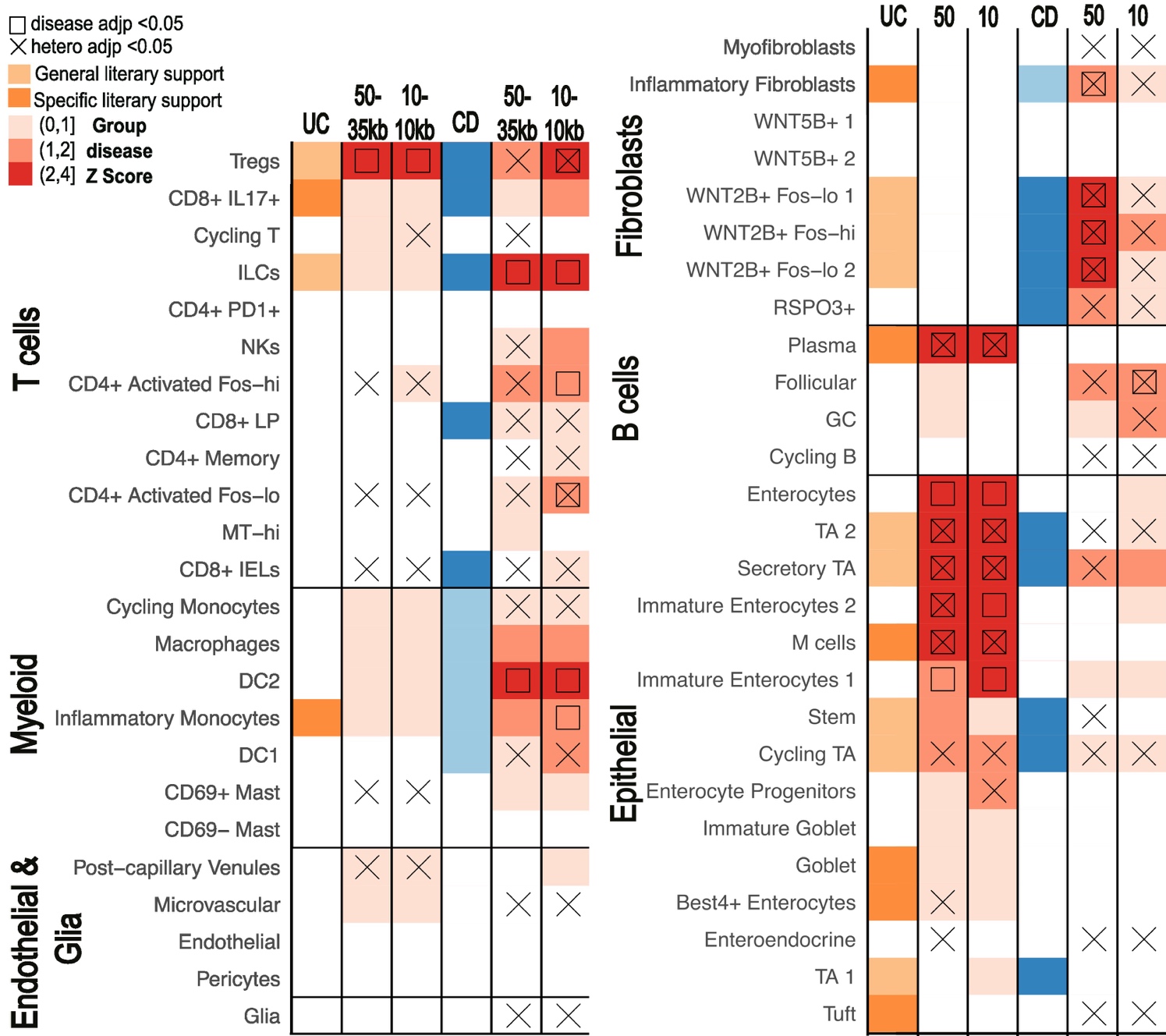
**Supplemental Figure 9. MAGMA window comparisons for ulcerative colitis and crohn's disease.** scDRS results of significant clusters for UC and CD using 50-35kb and 10-10kb windows. scDRS defines significant clusters with a group disease Z-score as shown in the gradient legend. Significance legend: *P<0.05, **P<0.01, ***P<0.001. UC=ulcerative colitis, CD=crohn’s disease.

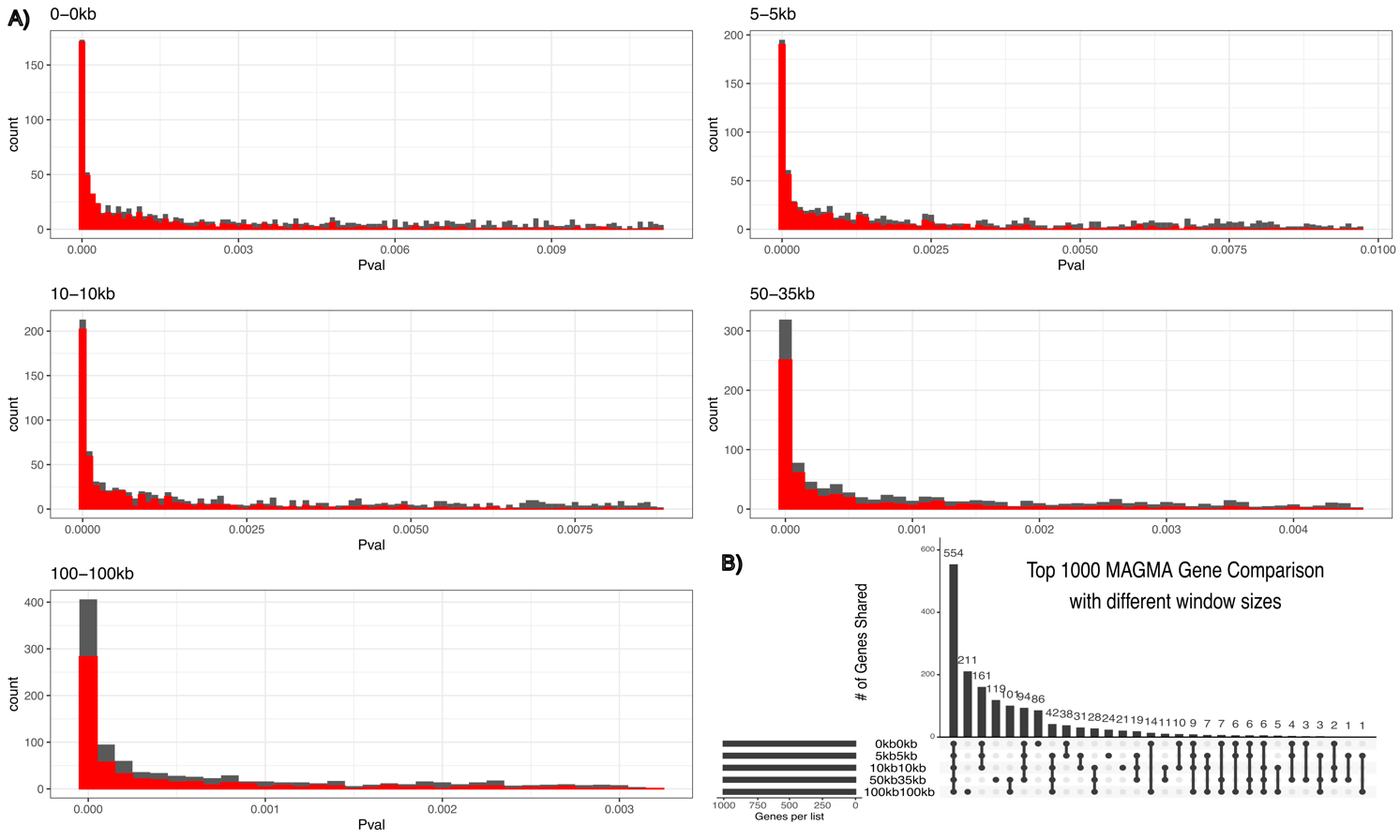
**Supplementary Figure 10.** The number of genes shared between the top 1000 genes deemed significant by MAGMA that also were found in scRNA-seq, with 5 different windows (0kb-0kb, 5kb-5kb, 10kb-10kb, 50kb-35kb, 100kb-100kb) **B)** The distribution of p-values of the the same top 1000 genes in B) with the distribution of the shared 554 genes highlighted in red.

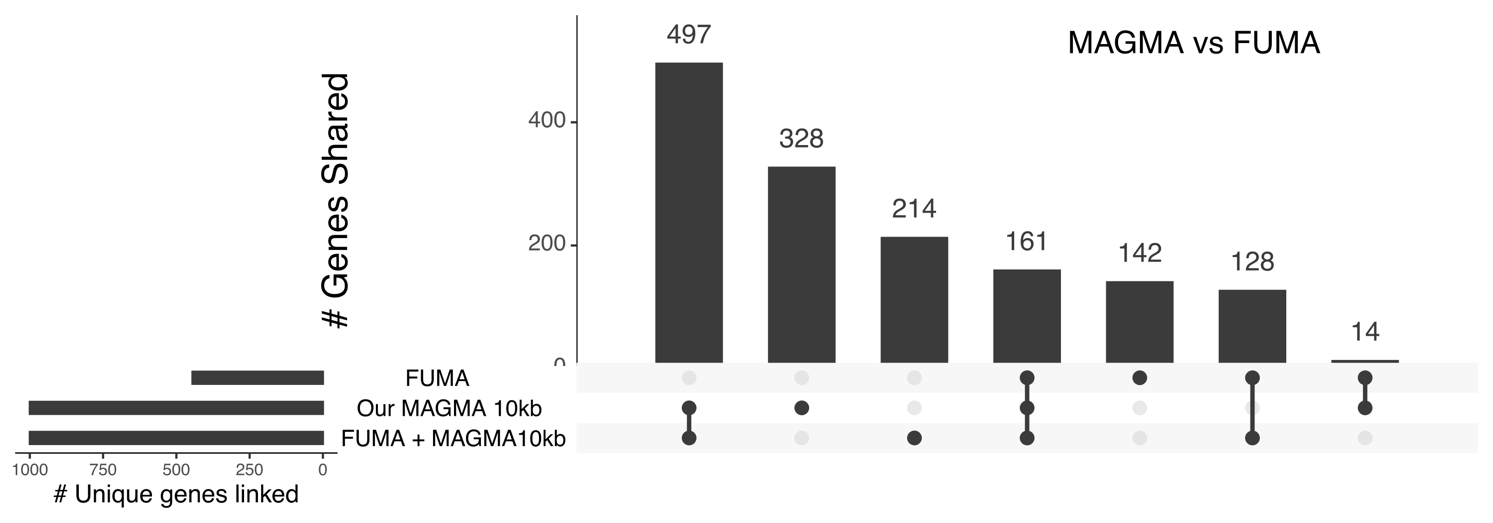
**Supplementary Figure 11:** UpSet plot of the mapped genes according to FUMA (FUMA), MAGMA run on FUMA’s final summary statistics (FUMA+MAGMA10kb), and MAGMA run on our final summary statistics (Our MAGMA 10kb)

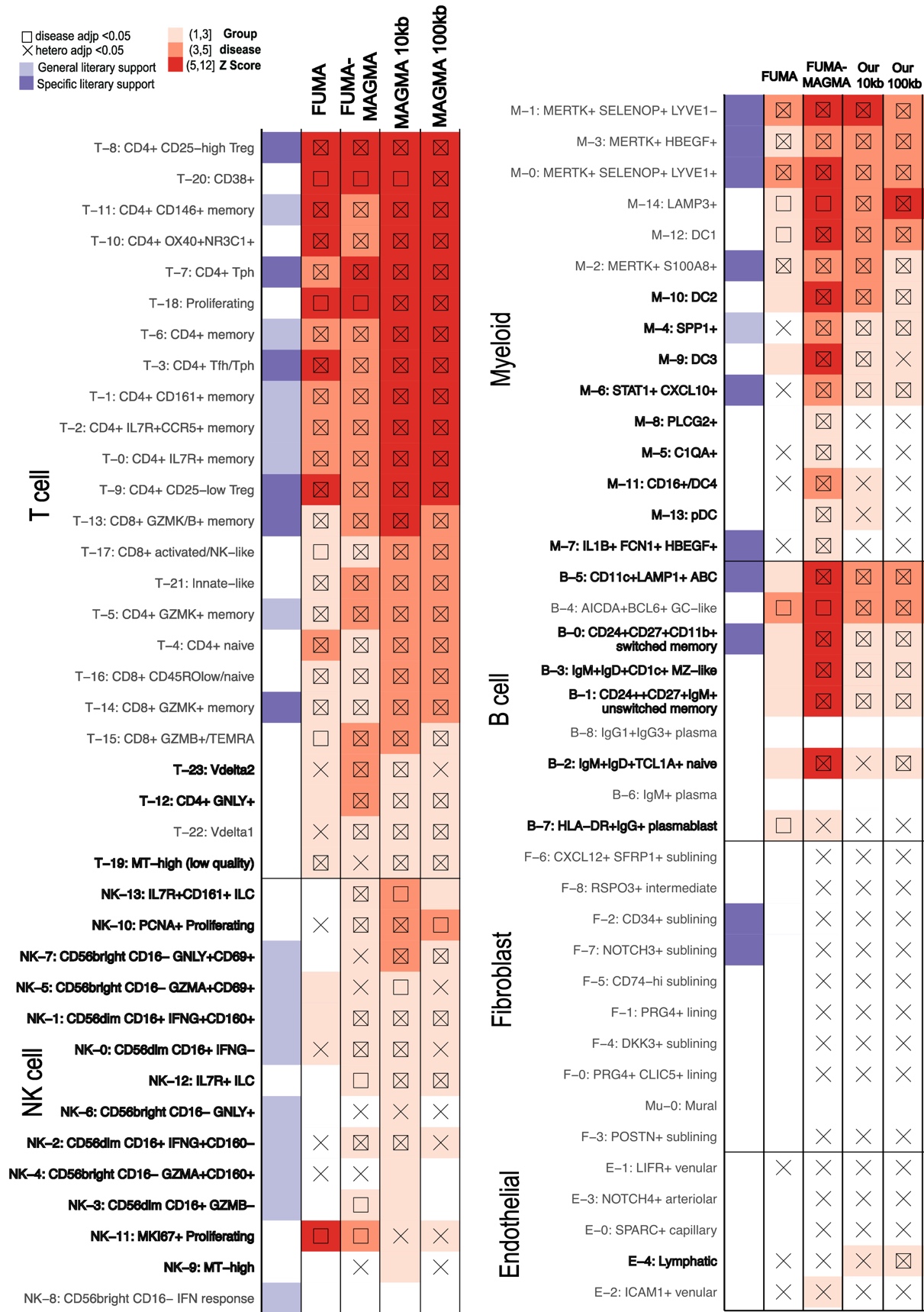

**Supplementary Figure 12:** scDRS results of significant clusters from rheumatoid arthritis calls using inputs from FUMA generated SNP analysis using MAGMA 10kb window mapping (FUMA-MAGMA), using FUMA based mapping including 10kb window, eQTL, and 3D chromatin interaction mapping (FUMA), and the 10kb/100kb MAGMA windows from Figure 5. Cell states with literary support are highlighted in purple. Cell states with differences in disease significance calls are bolded.
